# The “dark magic mushroom” co-produces amatoxins and psilocybin

**DOI:** 10.64898/2026.07.17.739257

**Authors:** Andrew R. Kunik, Michael W. Christopher, Benjamin Lemmond, Bryn T.M. Dentinger, Jason Slot, Timothy J. Garrett, Matthew E. Smith

**Affiliations:** Department of Plant Pathology, University of Florida; Gainesville, FL, USA; Department of Chemistry, University of Florida; Gainesville, FL, USA; Department of Pathology, Immunology and Laboratory Medicine, College of Medicine, University of Florida; Gainesville, FL, USA; Department of Plant and Microbial Biology, University of California, Berkeley; Berkeley, CA, USA; School of Biological Sciences and Natural History Museum of Utah, University of Utah; Salt Lake City, UT, USA; Department of Plant Pathology, The Ohio State University; Columbus, OH 43210, USA

## Abstract

The most famous chemicals produced by mushrooms are the psychedelic compound psilocybin from “magic mushrooms” and amatoxins from deadly poisonous mushrooms. These compounds are known to occur in multiple phylogenetically disjunct fungal lineages but have never been shown to co-occur within a single species. Here we show that the “dark magic mushroom” *Galerina indica* produces both psilocybin and amatoxins. Mass spectrometry revealed psilocybin and amatoxins in mushroom tissues, and genomic analyses identified corresponding biosynthetic genes. Phylogenetic analyses suggest that *G. indica* acquired psilocybin biosynthesis via horizontal gene transfer after amatoxin biosynthesis was already established, and that psilocybin biosynthesis was acquired twice independently within *Galerina*. Intriguingly, acquisition of psilocybin biosynthesis in *G. indica* may have coincided with reduced amatoxin potency. These findings reveal how horizontal gene transfer can combine powerful bioactive systems in a single species, potentially altering the ecological roles of both compound classes and the evolutionary fitness of the species.

## Introduction

Psilocybin and psilocin, collectively referred to as “psiloids,” are the principal alkaloids produced by psychedelic mushroom species (a.k.a. “magic mushrooms”) (*1, 2*). Psiloids and the mushrooms that produce them have long been valued in Mesoamerican Indigenous traditions and have attracted renewed scientific interest for their therapeutic potential in clinical psychiatry (*3–7*). By contrast, amatoxins are cyclic peptides present in the world’s deadliest mushrooms, including *Amanita phalloides* (the “death cap” mushroom). Amatoxins are responsible for most fatal mushroom poisonings through inhibition of RNA polymerase II and consequent organ failure (*8–10*). Although the effects of psiloids and amatoxins in humans are well documented, their natural roles in the environment remain unresolved, with anti-fungivore defense being the leading hypothesis for both compound classes (*10, 11*).

Although most specialized metabolite pathways in mushroom-forming fungi are inherited vertically, psiloid and amatoxin biosyntheses are canonical exceptions, with both pathways showing strong evidence of horizontal gene transfer between distantly related fungal species in different genera (*12–21*). In most psiloid-producing fungi, psiloid biosynthesis is encoded by a compact five-gene *Psi* cluster, whereas amatoxin biosynthesis depends on the *MSDIN/POPB* gene system (*20, 22–25*). Phylogenetic analyses of both systems show strong gene-tree/species-tree discordance, consistent with horizontal transfer (*15–20*). In the genus *Galerina*, deadly amatoxins are well documented in the *G. marginata* species complex (including *G. marginata, G. sulciceps, G. venenata* and others), which are colloquially referred to as “funeral bells” (*26, 27*). Until this study, amatoxins had only been confirmed from the *G. marginata* complex and no phylogenetically placed *Galerina* species had been reported to produce psilocybin or related psiloids (*28, 29*).

To search for novel amatoxin-containing species in the southeastern United States, we conducted exploratory chemical analyses of understudied *Galerina* species. Remarkably, we found that *Galerina indica,* which we designate the “dark magic mushroom,” co-produces psiloids and amatoxins, therefore dispelling the prevailing expectation that these two compound classes do not co-occur in the same species. We also discovered a second, distantly related *Galerina* species that produced psiloids but not amatoxins. We sequenced genomes to confirm the corresponding biosynthetic genes and examine their evolutionary relationships with homologs from other psiloid- and amatoxin-producing fungi. Our findings suggest that amatoxin biosynthesis was incorporated into the genus *Galerina* once, whereas psiloid biosynthesis was horizontally transferred in the genus at least twice: once into a non-toxic lineage, and once into an amatoxin-producing lineage, giving rise to the “dark magic mushroom” (Figure 1).

**Figure 1.**
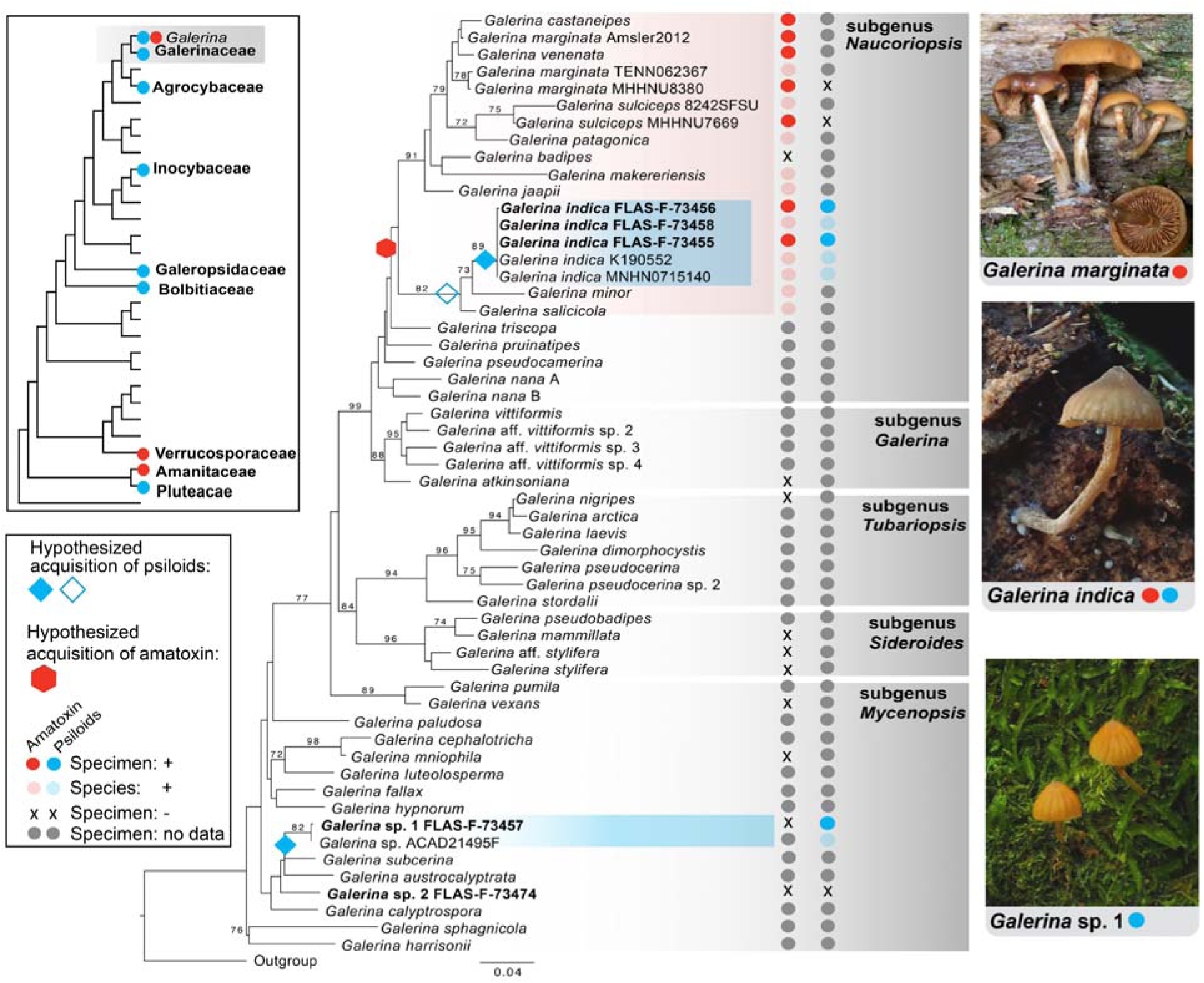
Organismal phylogenies of the genus *Galerina* and the order Agaricales. Box in upper left: Schematic showing the known distribution of psiloids and amatoxins in genera of Agaricales. Topology is based on a family-level, genome-based phylogeny in Niskanen et al. (*31*). Center: Maximum-likelihood phylogeny of the genus *Galerina* inferred from a concatenated three-locus alignment (ITS, 28S, *RPB2*). Red and blue icons placed over branches indicate hypothesized horizontal gene transfer acquisitions of the psiloid (blue) and amatoxin (red) biosynthesis pathways. Specimens with chemical evidence for amatoxins and/or psiloids (this study, Landry et al. [*27*]) are annotated to the right of the tree with red or blue circles, whereas black “X” symbols indicate a confirmed negative result. Half-shaded red or blue circles indicate inferred presence of compounds based on phylogenetic placement whereas grey circles indicate lack of data. Bootstrap support >70% is indicated on branches. Right: Photographs of representative species with the three observed bioactive chemical profiles are shown at right: *G. marginata* (top, amatoxins only), *G. indica* (middle, both amatoxins and psiloids), and *Galerina* sp. 1 (bottom, psiloids only) (Photos: A. Kunik).

## Results

### *Galerina* specimen collection and phylogenetic placement

We collected five *Galerina* specimens in the southeastern USA (Table S1). Phylogenetic analysis of concatenated ITS-28S- *RPB*2 sequences indicated that these specimens represent three distinct, species-level lineages (Figure 1 and Figure S1-S4 and Table S2). The first three specimens (FLAS-F-73455, FLAS-F- 73456, and FLAS-F-73458) found emerging from moss-covered palm trunks in forested wetlands (Florida, USA), were identified as *Galerina indica* (*30*). *Galerina indica* belongs to subgenus *Naucoriopsis*, which includes the deadly toxic *G. marginata* species complex (Figure 1) (*26, 27*).

The fourth specimen, *Galerina* sp. 1 (FLAS-F-73457), found emerging from a moss- covered dead pine (Mississippi, USA), could not be assigned to known species and was phylogenetically resolved in subgenus *Mycenopsis*. The fifth specimen, *Galerina* sp. 2 (FLAS-F- 73474), found emerging from moss-covered soil (Florida, USA), could not be assigned to a known species and was also phylogenetically placed within subgenus *Mycenopsis*.

### Chemical analysis

Higher-energy Collisional Dissociation (HCD) tandem mass spectrometry (MS/MS) analysis with authenticated standards showed that all analyzed collections of *G. indica* contained psiloids (psilocin and psilocybin) and two amatoxin compounds at *m/z* (a unitless value corresponding to the ratio of ion mass to the charge number) 903.367 and 887.370. The dominant peak at *m/z* 903.367 with retention time (RT) of 8.99 min was structurally characterized and found to be a novel amatoxin that we designate α-proamanitin. Its structure (Figure 2) is otherwise identical to α-amanitin, the most common of the deadly amatoxins, but lacks the hydroxyl group at the C-4 position of the proline residue. The stereochemistry shown for α-proamanitin is inferred from α-amanitin.

**Figure 2.**
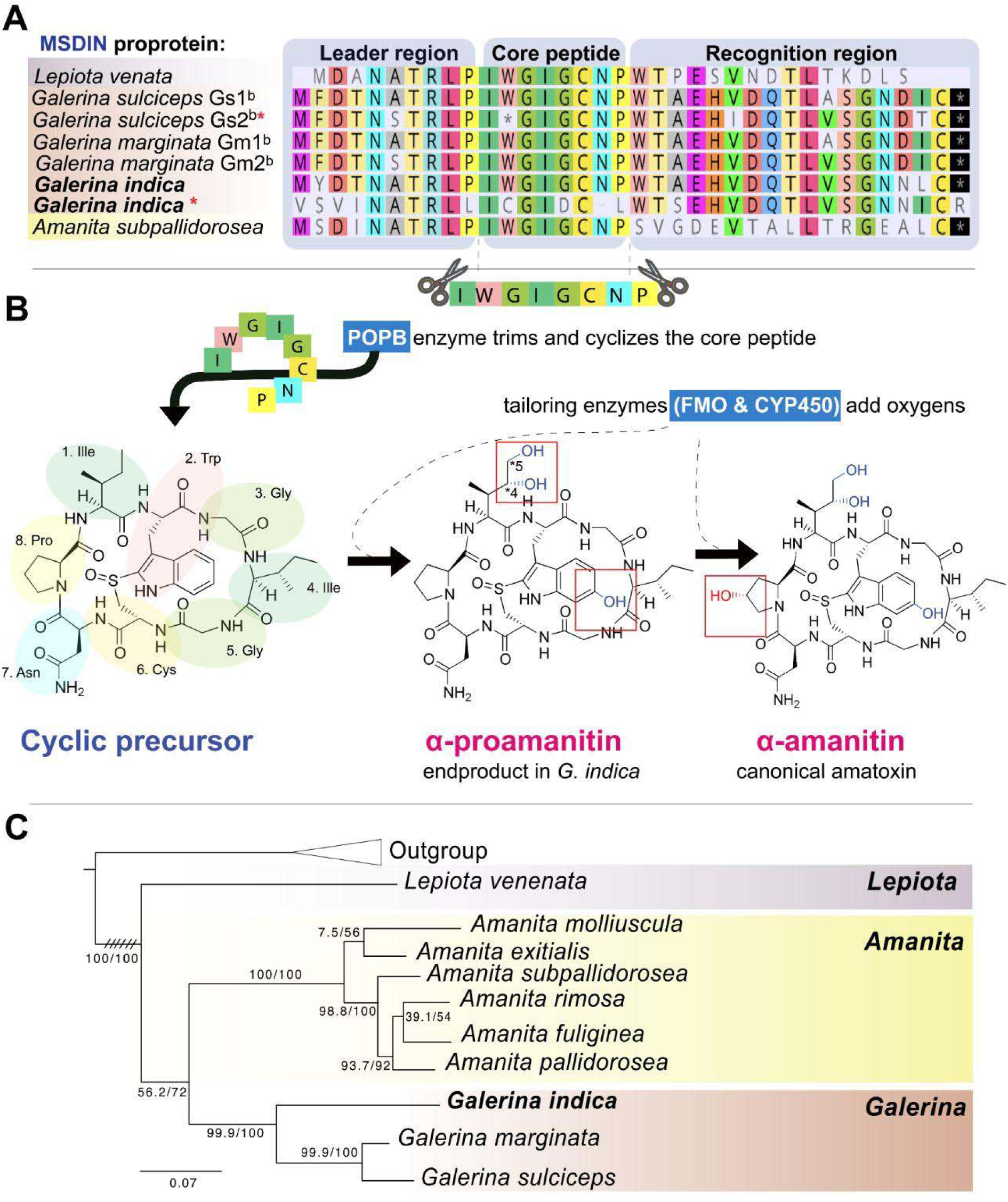
Amatoxin biosynthesis. **(A)** Alignment of MSDIN proproteins. Red asterisks denote putatively nonfunctional copies in *Galerina indica* and *G. sulciceps*. The nonfunctional *G. indica* copy contains a single-nucleotide deletion in the core region that causes a frameshift; the affected codon position is represented by a dash in the translated alignment at the position corresponding to the conserved “N” core residue. **(B)** Schematic showing the biosynthetic pathway that produces α-amanitin or α-proamanitin. First, the core peptide sequence is trimmed from the MSDIN proprotein. The core peptide is then cyclized, creating the cyclic precursor. Finally, tailoring enzymes hydroxylate isoleucine and tryptophan to create α-proamanitin in *G. indica*. Additional tailoring enzymes hydroxylate the proline to create the canonical amatoxin α-amanitin in *G. marginata* and most other amatoxin-containing taxa. (**C**) Maximum-likelihood phylogeny of POPB protein sequences. The tree is rooted with paralogous housekeeping POP proteins that are not involved in amatoxin biosynthesis, using sequences from related Agaricales mushrooms in the genera *Hypsizygus* and *Termitomyces*. Branches overlaid with three sets of double slashes indicate that the displayed branch length has been compressed to one-eighth of its original length.

Structural characterization of the ion at *m/z* 887.370 and RT 8.99 min supported a structure that matches α-proamanitin but lacks an additional hydroxyl group on the first isoleucine residue. MS/MS alone cannot resolve the position of this missing isoleucine hydroxyl, but it is likely either the C-4 or the C-5 position given the context of other known amatoxins (*10*). Notably, α-amanitin was not detected in any *G. indica* samples. All psiloid spectra and structural justification for both amatoxins are provided in (Supplementary Note and Figures S5- S15).

*Galerina* sp. 1 contained psilocin and psilocybin but no detectable amatoxins. *Galerina* sp. 2 contained no detectable levels of either class of these compounds. Raw mass spectrometry data from all analyzed specimens were deposited at MassIVE (accession number MSV000101888).

### Genome sequencing and assembly

Genomic DNA from *Galerina indica* (FLAS-F-73455) and *Galerina* sp. 1 (FLAS-F-73457) were sequenced using Illumina whole-genome shotgun sequencing, and *G. indica* was also sequenced with Oxford Nanopore long-read sequencing.

Genome assemblies were 61.65 Mb with an N50 of 90,626 bp and 98.2% BUSCO completeness for *G. indica*, and 63.2 Mb with an N50 of 2,415 bp and 68.1% BUSCO completeness for *Galerina* sp. 1. Assembly statistics and completeness metrics are provided in Table S3. Raw reads, assemblies, and gene annotations were deposited under BioProject PRJNA1468300.

### Identification and analysis of *Psi* genes

All five of the canonical *Psi* genes implicated in psiloid biosynthesis were identified in the genomes of *G. indica* and *Galerina* sp. 1. Gene order and orientation differed between them; *G. indica* had an order/orientation of *+PsiD+PsiM+PsiT2-PsiH-PsiK,* whereas *Galerina* sp. 1 had the order/orientation of *+PsiD- PsiH-PsiK+PsiM+PsiT2* (Figure 3 and Figure S16). These arrangements are evolutionarily informative because independent horizontal gene acquisition events are implicated by distinct *Psi* gene orders (*16, 17*).

**Figure 3.**
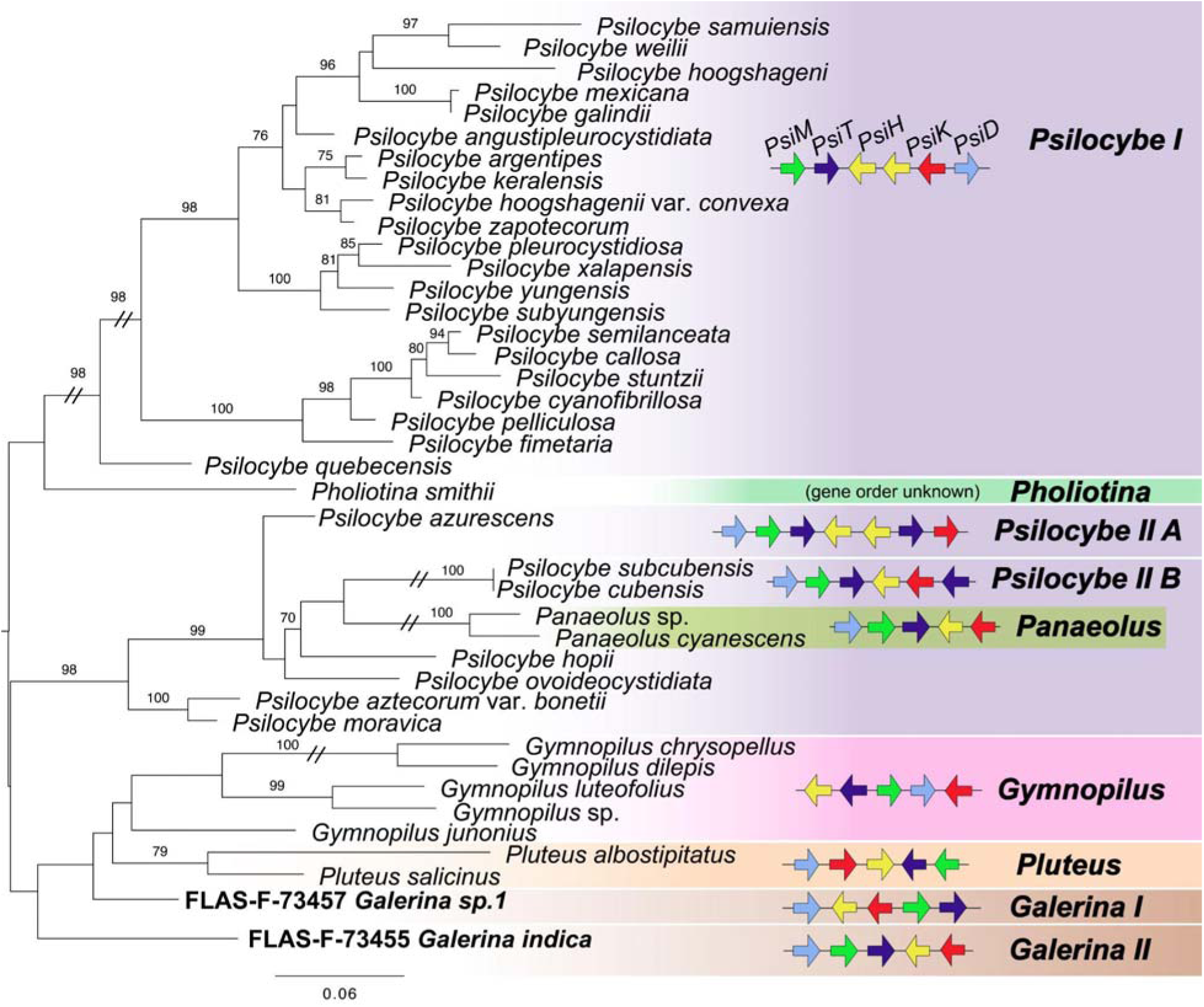
Psiloid biosynthetic gene phylogeny and gene-order. Maximum-likelihood protein phylogeny of PsiD. Where available, *Psi* gene order is provided in color-coded synteny diagrams. Each arrow represents a characterized gene in the biosynthetic pathway: *PsiM* (green), *PsiT* (dark blue), *PsiH* (yellow), *PsiK* (red), and *PsiD* (light blue). The phylogeny is midpoint rooted and bootstrap support >70% is indicated on branches. Branches overlaid with double-slash marks indicate that the branch length has been compressed by one-half.

We constructed protein phylogenies for four of the Psi proteins (PsiD, PsiH, PsiK, and PsiM) using our *Galerina* sequences together with homologs from previously characterized psiloid- producing taxa (including *Gymnopilus, Panaeolus, Pholiotina, Pluteus*, and *Psilocybe*) (Table S4). Midpoint-rooted phylogenies of the four Psi proteins show similar topologies, with *Galerina* sequences (*G. indica* FLAS-F-73455, *Galerina* sp. 1 FLAS-F-73457) not recovered as monophyletic in any phylogeny (Figure 3 and Figures S17-S19). Instead, *G. indica* and *Galerina* sp. 1 occurred on long, unsupported branches basal to or intermixed with *Pluteus* and *Gymnopilus*. Together, the differing *Psi* gene orders and the divergence between the two *Galerina* species in the organismal and Psi protein phylogenies are consistent with at least two independent acquisitions of the *Psi* cluster within *Galerina*. This interpretation is also consistent with the most parsimonious reconstruction of psiloid distribution on our *Galerina* organismal phylogeny (Figure 1): two acquisitions and no inferred losses, rather than one ancestral acquisition followed by approximately 12 inferred losses. The *Psi* biosynthetic gene cluster may have been acquired only in *G. indica* or may have been acquired earlier, in the ancestor of *G. indica, G. salicicola,* and *G. minor* (hollow blue diamond, Figure 1). However, more chemical and genomic data are needed from the rarely encountered species *G. salicicola* and *G. minor* before these two alternative scenarios can be adequately evaluated.

### Identification and analysis of amatoxin biosynthesis genes

Chemical analysis showed that *Galerina indica* contained amatoxins, so we searched its genome for core amatoxin-biosynthesis genes (*18, 24, 32*). Two *MSDIN* copies were detected: one inferred functional copy and one putatively non-functional copy (Figure 2). The functional copy occurred in a compact ∼6 kb cluster with *POPB* and *FMO1*, arranged *-POPB+FMO1-MSDIN*, matching the canonical amatoxin locus in other toxic *Galerina* species (Figure S20) (*19*). The putatively non-functional copy showed features of pseudogenization, including a mutation of the initiating methionine codon, resulting in M→V, substitutions in conserved prolines at predicted cleavage-sites (P→L) (*32*), and a single-base deletion in the core region that shifts the downstream reading frame.

The predicted protein sequences of both *MSDIN* gene copies were aligned with representative homologs from other known amatoxin-producing mushroom taxa (Figure 2 and Table S5). *Galerina sulciceps* and *G. marginata* also each contain two *MSDIN* gene copies (*25*). The inferred functional copy in *G. indica* contains the conserved core sequence “IWGIGCNP,” consistent with the amatoxin structures identified by our chemical analysis (e.g. α-proamanitin). In the amatoxin-producing taxa shown, MSDIN proteins contain the same “IWGIGCNP” core sequence as *G. indica*, but yield the canonical amatoxin α-amanitin as the pathway end product (*20, 24, 25, 33*).

A POPB protein phylogeny was constructed using the inferred *G. indica* amatoxin-cluster POPB protein and homologs from other amatoxin-producing taxa, including *Amanita* spp., *Lepiota venenata*, and members of the *G. marginata* complex (Figure 3 and Table S6). The three *Galerina* POPB protein sequences formed a strongly supported group, with *G. indica* on a long branch and sister to a clade containing *G. marginata* and *G. sulciceps*. This topology is concordant with the *Galerina* organismal phylogeny (Figure 1) and, together with the conserved amatoxin biosynthetic gene-cluster organization, supports a single origin of amatoxin genes within the genus *Galerina*. Together, these results support a scenario in which amatoxin production preceded psiloid production in *Galerina* subgenus *Naucoriopsis* (Figure 1). An alternative hypothesis requiring the loss of the *Psi* biosynthetic gene cluster from the ancestor of the *G. marginata* complex is less parsimonious.

## Discussion

Psychedelic and deadly poisonous mushrooms epitomize humanity’s enduring cultural fascination with fungi. Remarkably, the biosynthetic pathways responsible for psiloid and amatoxin production co-occur in *Galerina indica*. Horizontal gene transfer, although uncommon in eukaryotes, underlies this chemical combination and disrupts long-standing expectations about which fungal lineages produce which metabolites. In our chemical investigation of several obscure *Galerina* species, we expected to detect amatoxins, which are well documented in some members of the genus, but were surprised to find psiloids. These findings underscore the dynamic and unpredictable nature of chemical diversity among the vast number of understudied species in the fungal kingdom. Our results also have important public health implications. The blue pigmentation resulting from psilocin oligomers, often used to discriminate between co- habiting psychedelic and amatoxin-producing mushrooms, may only be useful in well-characterized or cultivated “magic mushroom” species that have a tradition of human use, such as *Psilocybe cubensis*. Indeed, we show that this blueing reaction can accompany amatoxins in some rare and uncharacterized species, as is the case of the “dark magic mushroom” *G. indica*.

Although several amatoxin variants have been characterized in previous research, α- amanitin is the principal and most widespread end product across amatoxin-producing fungi and is among the most potent amatoxins known (*20, 24, 25, 33*). *Galerina indica*, however, produces α-proamanitin, a close structural relative of α-amanitin that lacks C-4 hydroxylation of the proline residue (Figure 2). Because the “IWGIGCNP” MSDIN core motif in *G. indica* is identical to that of other α-amanitin-producing fungi, this difference reflects disruption of a downstream post-translational oxidation step rather than a change in the precursor peptide itself (Figure 2). Our ability to evaluate the candidate genes responsible for this difference in *G. indica* is limited because the enzymes mediating proline hydroxylation in amatoxin biosynthesis are not fully characterized. Luo et al. (*19*) showed that FMO1 and P450-29 enzymes are involved in post-translational oxidation generally in *G. marginata*, but they did not determine the precise site of the proline hydroxylation step. Although we identified an inferred FMO1 homolog in the *G. indica* amatoxin biosynthetic locus (Figure S20), the P450 gene family described by Luo et al. (*19*) in *G. marginata* was too divergent to confidently identify a direct P450-29 ortholog in *G. indica*. Therefore, the accumulation of α-proamanitin in *G. indica* may reflect lineage-specific differences in post-translational oxidation, raising the possibility that additional chemically distinct amatoxin intermediates or analogs remain to be discovered in *Galerina*.

Although the ecological functions of amatoxins and psiloids remain unresolved, both are hypothesized to mediate interactions with diverse organisms that consume fungi, such as vertebrates, mollusks, nematodes, and arthropods (*10, 11*). Because amatoxins inhibit eukaryotic RNA polymerase II, they are thought to function primarily as post-ingestive defenses against animal fungivores (*10, 34–36*). Psiloids may instead act behaviorally, reducing grazing or altering movement and oviposition in ways that could either indirectly reduce damage to mushrooms or directly enhance spore dispersal (*11*). The co-occurrence of both types of compounds in *G. indica* suggests a potential selective advantage in maintaining both pathways despite added biosynthetic investment (*11, 19, 37*). This pattern resembles “pyramidal” or “portfolio” defense strategies described in other systems, such as milkweed (*Asclepias* spp.) and pyramided Bt crops, in which distinct toxin combinations can synergistically reduce consumer performance and slow enemy counter-adaptation (*38–40*). Laboratory studies show that nematodes and fruit flies can evolve amatoxin resistance, suggesting that grazing pressure could drive counter-adaptation (*34, 41*). This may be especially relevant for nematodes, which consume a substantial fraction of fungal biomass in soil ecosystems (*42*). Psychedelic tryptamines are also toxic to nematodes, altering behavior and suppressing feeding (*34, 43–45*). In the environment where *G. indica* is found (e.g. decaying palms in hot, humid swamps in Florida) this two-pronged chemical strategy may broaden deterrence against various natural enemies while promoting spore dispersal by adapted mycovores.

However, the pyramidal defense hypothesis assumes that α-proamanitin provides sufficient bioactivity for ecological relevance. The potency of α-proamanitin is called into question when considering the structure-activity relationships among known amatoxins. While α-proamanitin possesses the 4-hydroxyisoleucine moiety necessary for sufficient bioavailability (*10, 46*), it lacks the hydroxyproline residue important for direct inhibition of animal RNA polymerase II. In vitro comparative studies have demonstrated that amatoxins lacking hydroxyproline exhibit weaker polymerase II inhibition than their hydroxyproline-containing analogs (*47*), and co-crystal structures show that hydroxyproline contributes directly to RNA polymerase II binding through hydrogen-bonding interactions (*48, 49*). Although amatoxins lacking hydroxyproline remain toxic, these observations suggest that α-proamanitin may be less bioactive than α-amanitin and therefore may be a less effective defense on its own against mycovores. Nevertheless, defensive value need not depend solely on intrinsic potency. If α- proamanitin is less metabolically costly to produce, poses less risk of autotoxicity to *G. indica* itself, or has effects enhanced by co-occurring psiloids, then it would still contribute to an economical, multi-compound defense strategy.

The broad prevalence of α-amanitin across amatoxin-producing fungi, together with the comparative rarity of α-proamanitin, suggests that α-amanitin is generally the evolutionarily favored end product. Our results indicate that α-amanitin-producing *Galerina* species share a common ancestor with *G. indica*, consistent with an ancestral α-amanitin pathway that was later altered in the *G. indica* lineage. Although neutral degeneration of the α-amanitin pathway cannot be excluded as a possibility, we hypothesize that the acquisition of psiloid production may have relaxed selection on proline hydroxylation if the newly acquired psiloid pathway partly supplanted the defensive advantage of α-amanitin (comparable to replacement of muscarine by psiloids in *Inocybe* [*50*]) or created a “pyramidal” stacking of two complementary defense systems. Alternatively, prior loss of α-amanitin production may have created selective conditions favoring retention of a horizontally acquired psiloid biosynthetic cluster, compensating for reduced amatoxin potency. In either scenario, *G. indica* brings together in a single species the biosynthetic systems associated with both “magic mushrooms” and the world’s deadliest poisonous mushrooms, providing a rare natural system for studying how horizontally acquired secondary-metabolite pathways impact organismal fitness.

## Methods

### Field Work and Morphological Observations

Fresh basidiomata of *Galerina* were photographed in situ and the substrate and habitat were recorded at the time of collection. Macroscopic photographs were obtained with an Olympus TG-7 camera (OM Digital Solutions, Tokyo, Japan). Hand sections were mounted in 3% KOH for microscopic study and observed with a Zeiss Axio Imager A2 compound microscope (Carl Zeiss, Oberkochen, Germany) under brightfield/DIC optics. Images of basidiospores were captured using an Axiocam 305 camera and their length and width measured in profile view with Zen Pro v. 3.1 software (Carl Zeiss, Oberkochen, Germany). Specimens were dried in a food dehydrator at 45 °C for 3 hours and accessioned at the University of Florida Herbarium (FLAS) and Garrett Herbarium at the University of Utah (UT). Collection metadata, microscopic observations, and field photographs were deposited in iNaturalist and MycoPortal and links are provided in Table S1.

### Ribosomal DNA Barcoding

DNA was extracted from fruiting bodies of voucher specimens using the rapid alkaline extraction method of Rennick et al. (*51*). Nuclear rDNA internal transcribed spacer (ITS1–5.8S–ITS2) and large subunit (28S) regions were targeted for amplification. The ITS region was amplified with primers ITS1F/ITS4, and 28S with LROR/LR5 (*52, 53*). PCR reactions were performed in 20 μL volumes containing 0.8 μL of each 10 μM primer, 1 μL template DNA, 0.8 μL of 1% BSA, 2 μL of 10× buffer, 0.48 μL of 10 μM dNTPs, 0.16 μL Taq polymerase (New England Biolabs, Ipswich, MA), and nuclease-free water (OmniPur®, MilliporeSigma). Cycling conditions were: 95 °C for 5 min; 35 cycles of 95 °C for 30 s, annealing at 58 °C for 30 s with a –0.1 °C touchdown per cycle for the first 10 cycles followed by annealing at 57 °C for 25 cycles, and extension at 72 °C for 1 min; final extension at 72 °C for 10 min.

PCR products were stained with SYBR Green I (Lonza Bioscience, Rockland, ME), visualized on 1.5% agarose gels, and Sanger sequenced with the same primers (Eurofins Genomics, Louisville, KY). Sequences were manually trimmed and then assembled in Geneious 2020.2.4 (Biomatters, Auckland, New Zealand) using the “De Novo Assembly” function with default settings.

### *Galerina* organismal phylogeny

Species placement and relationships within *Galerina* were evaluated using 28S, ITS, and *RPB2* phylogenies. Reference sequences for the diversity in the genus *Galerina* were obtained from the supplementary tables of Landry et al. (*27*), which included metadata on amatoxin presence/absence and subgenus placement (Table S2). Newly generated sequences from our *Galerina* vouchers were queried against the NCBI nucleotide database using BLAST, and the closest related sequences not included in Landry et al. (*27*) were added to the data set. ITS sequences from the *Galerina* genomes from He et al. (*21*) implicated in the amatoxin biosynthetic gene analysis were also included. *Gymnopilus spectabilis* PBM2471CUW was used as an outgroup. Sequences were aligned using MAFFT v7.520 with the --auto algorithm-selection parameter. Poorly aligned regions were trimmed using TrimAl v1.4.1 with the -automated1 parameter. The *RPB2* alignment was validated as intron-free and in-frame. The three locus alignments were concatenated into a single 56-taxon, 1,981-bp matrix.

Boundaries between ITS subregions (ITS1, 5.8S, ITS2) were identified with ITSx v1.1b (*54*), with detected coordinates mapped back to alignment columns. The concatenated matrix was partitioned into seven candidate blocks: ITS1, 5.8S, ITS2, 28S, and *RPB2* codon positions 1, 2, and 3. Substitution model selection and partition merging were performed simultaneously by ModelFinder Plus (-m MFP+MERGE) (*55*) in IQ-TREE v2.4.0 (*56*), with the best scheme chosen by BIC. Phylogenetic inference was performed in IQ-TREE under the selected partitioning scheme, with branch support estimated from 1,000 ultrafast bootstrap replicates (*57*) using NNI optimization (--bnni) and gene-site resampling ([*58*]; --sampling GENESITE). The best scheme collapsed the seven candidate partitions to three: ITS1+ITS2 (TVM+F+G4), 5.8S+28S+*RPB2* codon positions 1+2 (TN+F+I+G4), and *RPB2* codon position 3 (HKY+F+G4). In addition to the concatenated analysis, single-locus trees were inferred for ITS, 28S, and *RPB2* under the same IQ-TREE settings. Taxa in the tree were manually annotated to denote presence/absence of amatoxins based on previous analysis and on data collected here (Table S2) (*27*, *21*, *59*). All phylogenetic trees generated in this study were visualized and annotated in FigTree v1.4.4 (http://tree.bio.ed.ac.uk/software/figtree/).

### Untargeted chemical analysis

Dried basidioma tissue (1–2 mg; a mixture of lamellae, stipe, and pileus tissue) from each voucher specimen was weighed and placed into a 1.5 mL graduated natural microcentrifuge tube (Fisherbrand Premium Microcentrifuge Tubes, Fisher Scientific, Pittsburgh, PA, USA; Cat. No. 05-408-129), each containing 10-25 0.7 mm zirconia beads (BioSpec Products, Bartlesville, OK, USA; Cat. No. 11079107zx). Tubes were sealed with Parafilm and homogenized in a Fisher Bead Mill 24 homogenizer (Fisher Scientific, Pittsburgh, PA, USA) for 30 s. Following homogenization, 1 mL of methanol was added to each tube, and samples were subjected to sonication (10 min, cold ultrasonic bath) to enhance metabolite extraction. Extracts were centrifuged in a Sorvall ST8R centrifuge (Thermo Fisher Scientific, Waltham, MA, USA) for 10 min at 20,000 × g at 4 °C. The resulting supernatant was collected, and 500 uL of the methanol extract was transferred into a new tube. Solvent was evaporated under a stream of nitrogen using an Organomation OA-HEAT evaporator (Organomation Associates, Inc., Berlin, MA, USA) on no heat mode for approximately 1 h until all residual methanol had dissipated. Dried residues were solubilized in 50 µL of water containing 0.1% formic acid. Solubilized extracts were centrifuged again under the same conditions, and 20 µL of the supernatant was transferred to an autosampler tube for subsequent analysis. Analytes were injected into an Avantor ACE Excel C18 column (100 × 2.1 mm, 2 μm) fitted with a HALO 90 Å C8 (5 × 2.1 mm, 2.7 μm) guard column. Mobile phase A was 0.1% water formic acid and mobile phase B was acetonitrile. The gradient started at 100% A for 3 min before linearly increasing to 80% B over the following 10 min. After holding 80% B for 3 min, the gradient was rapidly brought back to starting conditions to reequilibrate. Mass analysis was performed on a ThermoScientific Q Exactive Orbitrap mass spectrometer. Because some specimens were suspected to contain amatoxins or related bicyclic peptide toxins, spectra were initially screened for the formula masses of known amatoxins, phallotoxins, and related compounds (*10*). Psilocybin and psilocin were also among the most prominent features in several samples, prompting subsequent targeted analysis.

### Targeted chemical analysis of psiloids and amatoxin structural elucidation

The entire chemical analysis, including tissue resampling, was repeated using the same *Galerina* voucher specimens, together with analytical grade standards and two additional FLAS fungarium specimens with known metabolite profiles as controls: *Amanita suballiacea* (FLAS-F-60862), a species well known to contain amatoxins but not psilocybin or psilocin; and *Fomitopsis spraguei* (iNaturalist #228453130, FLAS), a polypore species expected to be devoid of psilocybin, psilocin, and amatoxins (Table S1). The control results were as expected and are not discussed further, and the raw mass spectra of all samples analyzed as a part of this study were deposited in MassIVE under the accession number MSV000101888. Analytical standards were purchased as follows: 4-hydroxy-*N,N*-dimethyltryptamine (psilocin; CAS 520-53-6) was purchased from Cayman Chemical (Ann Arbor, MI, USA) as a 1 mg/mL solution in acetonitrile, batch 0704075. γ-Amanitin (CAS 21150-23-2; 1 mg) was purchased from Apollo Scientific Ltd. (Denton, Manchester, UK), batch AS524140.

Targeted Higher-energy Collisional Dissociation (HCD) MS/MS of known amatoxins and of unknown amatoxins on *G. indica*, were performed. α-, and γ-Amanitin were used to generate reference spectra, which helped decipher product ion identities and localize where new amanitin species differed. α-Amanitin was sourced from *Amanita suballiacea* extract, a species known to produce α-amanitin through previous analysis and through our own exact mass matching and MS/MS spectral interpretation (*60, 61*).

### Genomic DNA Extraction

For genome sequencing of *Galerina indica* FLAS-F-73455, dikaryotic mycelium was established in axenic culture and grown on Difco™ malt extract agar (Becton Dickinson, Franklin Lakes, NJ; Cat. No. 211220) overlaid with dialysis paper. After approximately three weeks of growth, fresh mycelium was harvested from the dialysis paper and divided among four 2 mL tubes, each containing 40–100 mg of fresh tissue. Samples were frozen and macerated with a sterile pestle mounted to a drill press. DNA was extracted using the QIAGEN DNeasy PowerSoil Pro Kit ( Hilden, Germany; Cat. No. 47016), pooled across extractions, and further purified using the QIAGEN DNeasy PowerClean CleanUp Kit.

To obtain DNA for genome sequencing of *Galerina* sp. 1 (FLAS-F-73457) , 4-10 mg of dried fruiting body tissue was transferred to a 2 mL tube and moistened with cold lysis buffer from the E.Z.N.A.® HP Plant DNA Kit (Omega Bio-Tek, Norcross, GA, USA). The tissue was then macerated with a sterile pestle mounted to a drill press. Additional lysis buffer buffer was then added, and DNA was extracted using the E.Z.N.A.® HP Plant DNA Kit following the manufacturer’s protocol, except that 10 μL β-mercaptoethanol and 2 μL RNase were added to the lysis buffer and the lysate was incubated overnight at 55 °C.

### Illumina Sequencing, Processing, and Genome Assembly

The extracted DNA was submitted to the University of Florida Interdisciplinary Center for Biotechnology Research for standard Illumina library preparation using the NEBNext® kit and sequenced across two full lanes on a MiSeq i100 using the 25M reagent kit (2 × 300 bp; 15 Gb). Raw reads were trimmed of adapters filtered by quality and read length (<75bp) using AAFTF version 0.4.1 (*62*) with the bbduk method, followed by removal of PhiX and contaminant sequences using AAFTF filter. All trimmed and filtered paired-end libraries were provided as separate inputs to SPAdes version 4.0.0 (*63*) in isolate mode with k-mer values of 21, 33, 55, and 77. Contigs shorter than 500 bp were removed from the raw assembly. Assemblies were screened for foreign sequence contamination using NCBI Foreign Contamination Screen (FCS-GX) version 0.4.0 (*64*) using -- tax-id 109632 (*Galerina*) and the full GX database, and identified contaminant sequences were removed prior to downstream processing. The filtered assembly was then polished using POLCA, part of the MaSuRCA version 4.0.9 toolkit (*65*), with all six read libraries mapped back to the assembly to correct single-base errors. Reference-guided scaffolding was performed using RagTag version 2.0.1 (*66*) with the publicly available genome assembly of *Galerina marginata* GCA_023014335.1 as a reference. Assembly completeness was assessed using both BUSCO version 5.3.0 (*67*) with the agaricales_odb10 lineage dataset (n=3,870) and Compleasm version 0.2.7 (*68*) with the agaricales_odb12 lineage dataset (n=3,372). Raw reads and assemblies have been deposited in GenBank under BioProject PRJNA1468300.

### Oxford Nanopore Genome Sequencing and Assembly

A dikaryotic genomic DNA isolate of *Galerina indica* (FLAS-F-73455) was selected for additional hybrid long-read and short-read genome sequencing. Long-read libraries were prepared using the Oxford Nanopore Ligation Sequencing Kit V14 (SQK-LSK114) with enzyme E8.2.1 (Oxford Nanopore Technologies) and the Long Fragment Buffer. Libraries were sequenced on a PromethION R10.4.1 flow cell at the University of Florida Interdisciplinary Center for Biotechnology Research (ICBR). Base calling was performed with Dorado v1.3.0 using the super-accuracy model dna_r10.4.1_e8.2_400bps_sup@v5.2.0 on an NVIDIA B200 GPU. Oxford Nanopore reads were converted from unaligned BAM to FASTQ format using SAMtools v1.20 (*69*). Read quality and length distributions were assessed with NanoPlot v1.42.0 (*70*) and SeqKit v2.4.0 (*71, 72*). Reads were filtered with Filtlong v0.3.1 (https://github.com/rrwick/Filtlong) using --min_length 1000 -- keep_percent 95 --target_bases 8000000000 to retain the highest-quality 8 Gb of reads (∼74× diploid coverage / ∼147× per haplotype on the estimated 54.4 Mb haploid genome). Illumina reads to be used for polishing were quality-trimmed and adapter-clipped with fastp v0.23.4 (*73*) with parameters --detect_adapter_for_pe --qualified_quality_phred 20 --length_required 75 -- cut_tail --cut_tail_window_size 4 --cut_tail_mean_quality 20.

K-mer analysis was performed on trimmed Illumina reads using Jellyfish v2.3.0 (*74*) with k=21, -C (canonical k-mers). The resulting k-mer histogram was analyzed with GenomeScope2 v2.0 (*75*) under both haploid (-p 1) and diploid (-p 2) models to assess ploidy. The diploid model fit substantially better (97.6% vs. 88.7%), consistent with the expected dikaryotic state of the sample. Long-read de novo assembly was performed using Flye v2.9.3 (*76*) with parameters -- nano-hq --keep-haplotypes --min-overlap 5000 --genome-size 110m. Trimmed Illumina reads were aligned to the Flye assembly using BWA-MEM v0.7.17 (*77*), reporting all alignments (-a flag) for haplotype-aware polishing. Aligned reads were sorted and indexed with SAMtools v1.20.

Polypolish v0.5.0 (*78*) was applied in two steps: first, alignments were filtered by insert size using the bundled polypolish_insert_filter.py script; then polypolish polish corrected positions where short-read alignments unambiguously disagreed with the assembly. Subsequently, Pilon v1.24 (*79*) was run in diploid mode (--diploid --changes --vcf) on a fresh BWA-MEM alignment to the Polypolish output.

To produce a haploid primary assembly, purge_dups v1.0.1 (*80*) was applied to the polished assembly. Filtered Oxford Nanopore reads were aligned back to the assembly with minimap2 v2.28 (-x map-ont). Coverage cutoffs (5, 53, 87, 105, 175, 315) were determined automatically by calcuts, which identified two coverage peaks corresponding to haplotype-specific (∼70×) and homozygous/shared (∼150×) contigs. Haplotigs were identified through self-alignment with minimap2 (-x asm5 -DP) and removed with the purge_dups algorithm (*81*). The final primary assembly was screened for foreign contamination using NCBI FCS-GX v0.5.4 (*64*) against the GX-DB reference database (build date 2023-01-24) with --tax-id 109632 (*Galerina*). Assembly completeness was evaluated using compleasm v0.2.7 (*68*) and BUSCO v5.8.3 (*67*) with both agaricales_odb10 and basidiomycota_odb10 lineage datasets. Raw reads and assemblies have been deposited in GenBank under BioProject PRJNA1468300.

### Gene prediction and annotation

Protein-coding gene models were predicted using funannotate predict v1.8.17 (*82*), which integrates *ab initio* predictions with homology-based evidence via EVidenceModeler (*83*). Four *ab initio* gene predictors were employed: AUGUSTUS v3.5.0 (*84*), GeneMark-ES v4.73 in self-training mode (*85*), SNAP (*86*), and GlimmerHMM (*87*).

AUGUSTUS, SNAP, and GlimmerHMM were trained using conserved single-copy orthologs identified by BUSCO v2 (*88*) against the Dikarya lineage, with *Coprinopsis cinerea* parameters serving as the AUGUSTUS seed species. AUGUSTUS parameters were further refined using optimize_augustus.pl. Protein homology evidence was provided from two closely related reference genomes within Hymenogastraceae: *Galerina marginata* (GenBank GCA_000697645.1; [*89*]) and *Psilocybe cubensis strain* MGC- MH-2018 (RefSeq GCF_017499595.1; [*90*]). Reference proteins were aligned to the masked genome with DIAMOND v2.1.10 (*91*) followed by spliced alignment with Exonerate v2.4.0 (*92*). EvidenceModeler then integrated all *ab initio* predictions with protein alignments to produce weighted consensus gene models. Maximum intron length was set to 3,000 bp. Gene models shorter than 50 amino acids or with significant sequence homology to known transposable element proteins were removed. Transfer RNA genes were predicted using tRNAscan-SE v2.0.12 (*93*).

Predicted protein-coding genes were functionally annotated by integrating multiple evidence sources. Protein domains were identified using InterProScan v5.72-103.0 (*94*) against member databases including Pfam, SMART, PANTHER, and PRINTS. Orthology-based functional assignments, including Gene Ontology (GO) terms, KEGG pathway annotations, and COG categories, were obtained using EggNOG-mapper v2.1.12 (*95*) in DIAMOND search mode against the EggNOG 5.0 database (*96*). Biosynthetic gene clusters were identified using antiSMASH v7.0 (*97*) in fungal taxon mode, with ClusterBlast comparison to the MIBiG reference database (*98*), subcluster detection, Active Site Finder, Pfam-to-GO annotation, and secondary metabolism gene phylogenetic tree construction enabled. All functional annotation sources were integrated into the final gene models using funannotate annotate. Annotation completeness was evaluated using BUSCO v6.0.0 (*67*) in proteins mode against the basidiomycota_odb10 reference set.

### Identification of Psiloid Biosynthesis Genes from *Galerina* Genomes

Psiloid biosynthetic gene cluster detection was performed on each genome assembly using gene prediction followed by homology-based searches largely following the methods of (*16*). Gene prediction was first carried out using funannotate v1.8.17 (*82*) as described above. Predicted protein sequences were then searched for homologs of the four core psiloid pathway genes—PsiD (tryptophan decarboxylase; UniProt P0DPA6), PsiK (kinase; P0DPA8), PsiM (methyltransferase; P0DPA9), and PsiH (cytochrome P450 monooxygenase; P0DPA7)—from the publicly available *Psilocybe cubensis* reference assembly (*90*) using BLASTp (e-value threshold 1e-5). The top three hits for each reference gene were retained, and putative Psi gene clusters were identified computationally by examining the gene numbers assigned by Augustus: any group of two or more hits in which each gene was within two predicted gene positions of at least one other gene in the group was designated a putative cluster. Synteny visualization was performed using clinker v0.0.32 (*99*) with a minimum identity threshold of 30% for gene linking.

### Psiloid biosynthesis gene phylogeny

Psiloid biosynthetic gene sequences for phylogenetic analysis were extracted from the Augustus gene predictions of 57 previously published fungal genomes (52 *Psilocybe* spp., 2 *Gymnopilus* spp., 2 *Pluteus* spp. and 1 *Panaeolus* sp.) (Table S4). First, Augustus GFF annotation files containing predicted genes with embedded protein translations were downloaded from the Dryad Digital Repository (doi:10.5061/dryad.tmpg4f52s) for all genomes (*16*). Protein sequences were computationally extracted from GFF comment fields using custom Python scripts and compiled into a searchable BLAST database with makeblastdb (BLAST+ v2.15.0; [*100*]). To identify psiloid pathway homologs, reference protein sequences of the four core biosynthetic genes from the *Psilocybe cubensis* reference genome (*90*) were used as BLASTP queries with an e-value threshold of 1×10 and a maximum of 200 target sequences per query.

To retrieve the *Psi* genes from four recently sequenced genomes from Liu et al. (*17*) (NCBI BioProject accessions PRJNA1046270 and PRJNA1223616; genome assemblies *Gymnopilus sp.* GCA_051998105.1, *Gymnopilus luteofolius* GCA_051998225.1, *Panaeolus sp*. GCA_051997735.1, and *Psilocybe keralensis* GCA_051997275.1), psiloid biosynthetic genes were identified using a targeted gene prediction approach, as annotated protein predictions were not publicly available. Genome assemblies were downloaded from NCBI and nucleotide BLAST databases were constructed using makeblastdb. The same *P. cubensis* reference proteins were used as queries in tblastn searches (e-value ≤ 1×10 ² ) to identify candidate gene loci. Genomic regions flanking the top BLAST hit for each gene (±20 kb) were extracted using blastdbcmd, and spliced gene models were predicted using Exonerate v2.4.0 (*92*) in protein2genome mode with the --ryo "%tcs\n" flag to output predicted coding sequences. Predicted coding sequences were translated to protein using custom Python scripts, yielding four sequences per gene (one per genome, 16 total sequences).

Individual protein phylogenies were constructed for the psiloid biosynthetic genes PsiD, PsiH, PsiK, and PsiM. For each gene, multiple sequence alignment was performed using MAFFT v7.520 with the L-INS-i algorithm (--localpair --maxiterate 1000). Alignments were trimmed using TrimAl v1.4.1 with the -automated1 heuristic. The four alignments had the following lengths: PsiD, 395 amino acid positions; PsiH, 472; PsiK, 351; and PsiM, 294. The best-fit amino acid substitution model was selected independently for each gene partition in RAxML using the PROTGAMMAAUTO criterion, which selected JTT+Γ for *PsiD* and *PsiK*, JTTDCMUT+Γ for *PsiH*, and CPREV+Γ for *PsiM*. The trees were midpointed rooted in FigTree v1.4.4 (http://tree.bio.ed.ac.uk/software/figtree/). Synteny diagrams of Psi clusters were adapted from gene order and orientation data from this study and from Bradshaw et al. (*16*), Liu et al. (*17*), and Awan et al. (*23*).

### Identification of amatoxin biosynthesis genes from *Galerina* genomes

Amatoxin pathway genes were identified using a comprehensive dual-BLAST approach targeting three known biosynthetic enzymes: MSDIN (amatoxin precursor peptide), POPB (prolyl oligopeptidase), and FMO1 (flavin monooxygenase) following methodologies of Luo et al. (*19*) and Drott et al. (*101*). Reference sequences for MSDIN genes were compiled from a sample of amatoxin-producing species including *Galerina marginata* (GenBank: MN272413, MN272414), *G. sulciceps* (MN272417, MN272418), and *Lepiota venenata* (MN272421, MN272422), along with nucleotide coding sequences. Reference sequences for FMO1 (Galma1_104945), and POPB (MN272416, MN272420) were obtained from the JGI *Galerina marginata* v1.0 genome (Galma1) (*21*, *89*). Gene prediction was first carried out using funannotate v1.8.17 as described above. For maximum detection sensitivity, both protein-based searches (BLASTP against predicted proteins, e-value ≤ 1×10 ) and nucleotide-based searches (BLASTN and TBLASTN against the genome assembly, e-value ≤ 10 for MSDIN, ≤ 1×10 ³ for other genes) were performed using NCBI BLAST+ v2.15.0 (*100*). Synteny visualization was performed using clinker v0.0.32 (*99*) with a minimum identity threshold of 30% for gene linking, and *Galerina* genomes used from the literature that were without publicly available protein predictions were annotated in-house using funannotate v1.8.17.

### Amatoxin biosynthesis gene (POPB) phylogeny

Additional POPB protein sequences from other amatoxin containing species were obtained from He et al. (*21*) Table S6). Phylogenetic analysis largely followed methods in He et al. (*21*). Multiple sequence alignment of the 12 translated sequences was performed using MAFFT v7.520 with the --auto parameter, and poorly aligned or divergent regions were subsequently trimmed using trimAl v1.4.1 with the automated1 heuristic, yielding a final alignment of 728 amino acid sites. Maximum likelihood phylogenetic analysis was conducted in IQ-TREE2 v2.2.2.7, with the best-fit substitution model selected by ModelFinder Pro under the Bayesian Information Criterion. The Q.plant+G4 model was selected, incorporating a discrete Gamma distribution with four rate categories (α = 0.865). Branch support was evaluated using 1,000 ultrafast bootstrap replicates and 1,000 SH-aLRT tests. The tree was rooted using the POP sequences of *Hypsizygus marmoreus* (NCBI LUEZ02000233) and *Termitomyces sp.* (NCBI KQ412502) as outgroups.

## Data and materials availability

Genome assemblies and raw sequence reads have been deposited in GenBank under BioProject PRJNA1468300. Raw mass spectra of all analyzed specimens have been deposited at MassIVE under the accession number MSV000101888. Additional supporting data, such as alignments, are available at https://osf.io/jegf3/.

## Acknowledgments

This project was completed as part of an extended undergraduate research project by A. Kunik. The authors thank C. Dervinis and M. Kirst for troubleshooting molecular biology protocols, T. Kunik and C. B. Willis for help with processing specimen vouchers, Z. C. Smith for advice on genome assembly, A. Bradshaw for discussion of phylogenetic methods, and J.P. Cook and S. Ostuni (Florida Mycological Society) for help with logistical support and accessing Florida collecting sites. Psilocybin and psilocin possession is permitted under DEA licenses RD0672609 to B. Dentinger and RS0648913 to J. Slot.

## Author Contributions

Conceptualization: A.R.K. and M.E.S.; Methodology: A.R.K., M.W.C., and B.L.; Visualization: A.R.K., B.L., and M.W.C.; Funding acquisition: M.E.S. and T.J.G.; Project administration: M.E.S., J.S., and B.T.M.D.; Supervision: M.E.S., T.J.G., J.S., and B.T.M.D.; Writing – original draft: A.R.K., M.W.C., and M.E.S.; and Writing – revision & editing: A.R.K., M.W.C., B.L., J.S., B.T.M.D., and M.E.S.

