## Supplementary Materials for "The “dark magic mushroom” co-produces amatoxins and psilocybin"

**The PDF file includes:**

Supplementary Note
Figures S1 to S20
Tables S1 to S6
SI References

**Supplementary Note**

***Galerina indica* amatoxin structural characterization.** Initially, the feature at m/z 903.367, 8.6 min from *Galerina indica* (FLAS-F-73455), was assumed to be γ-amanitin, based on matching exact mass and isotope pattern. However, a γ-amanitin standard, run under the same conditions, eluted earlier at 8.46 min and had a different MS/MS spectrum (Figures S5 and S6). The most obvious difference in the unknown MS/MS spectrum compared to the γ-amanitin MS/MS spectrum was the presence of m/z 70.066 and 86.097, and the lack of a dominant peak at m/z 86.061 (Figure S6). α-Amanitin MS/MS spectra from *Amanita suballiacea* (FLAS-F-60862) were then used as another point of reference to compare the spectra and help deduce where the isomeric difference occurs.

The molecular formulas assigned to m/z 70.066 and 86.097 were $C_4H_8N^+$ and $C_5H_{12}N^+$, respectively. Interestingly, the dominant m/z peak in this range in α- and γ-amanitin MS/MS spectra is m/z 86.060 ($C_4H_8NO^+$), which corresponds to the addition of an oxygen to m/z 70.066 (Figure S7). Therefore, the m/z 70.066 signal provided us with the smallest substructure of the unknown compound that lacks an oxygen relative to γ-amanitin. The simplest explanation for where this oxygen was removed from in the unknown is on the proline, as the molecular formula matched and proline is one of the few variable sites across amanitin chemical species. To verify this, we found other product ions that should contain the proline and additional adjacent amino acid residues. Larger product ions that contain the proline agree with the initial localization of oxygen removal (Figures S8 to S12).

To finalize the unknown chemical structure, the localization of the missing hydroxyproline oxygen had to be assigned, and all other variable oxygens had to be consistent against γ-amanitin. Fortunately, the product ions shown in Figure S12 provide conclusive localization of the oxygen to the 5-position of hydroxyisoleucine, making it a dihydroxyisoleucine like what is seen for α-amanitin. m/z 243.133 is a unique product ion that contains both the variably hydroxylated isoleucine and proline, and as expected, is present in both γ-amanitin and the unknown amanitin. α-Amanitin does not have the m/z 243.133 product ion because it is shifted by an oxygen to m/z 259.128.

The more minor amanitin feature found in *Galerina indica* corresponded to a loss of one oxygen compared to pro-α-amanitin and had an m/z 887.370 at RT of 9.0 min. MS/MS of the feature showed the oxygen had been removed from the dihydroxylated isoleucine to form a singularly hydroxylated isoleucine. m/z 227.140 and 307.130 localized the missing oxygen to the isoleucine/proline portion of the peptide, and the presence of a product ion at m/z 70.066 gave high confidence that the proline remained non-hydroxylated (Figure S13). The exact position of the missing hydroxy group on isoleucine cannot be determined from MS/MS alone; however, we can confidently say the isoleucine is mono-hydroxylated.

### Figures

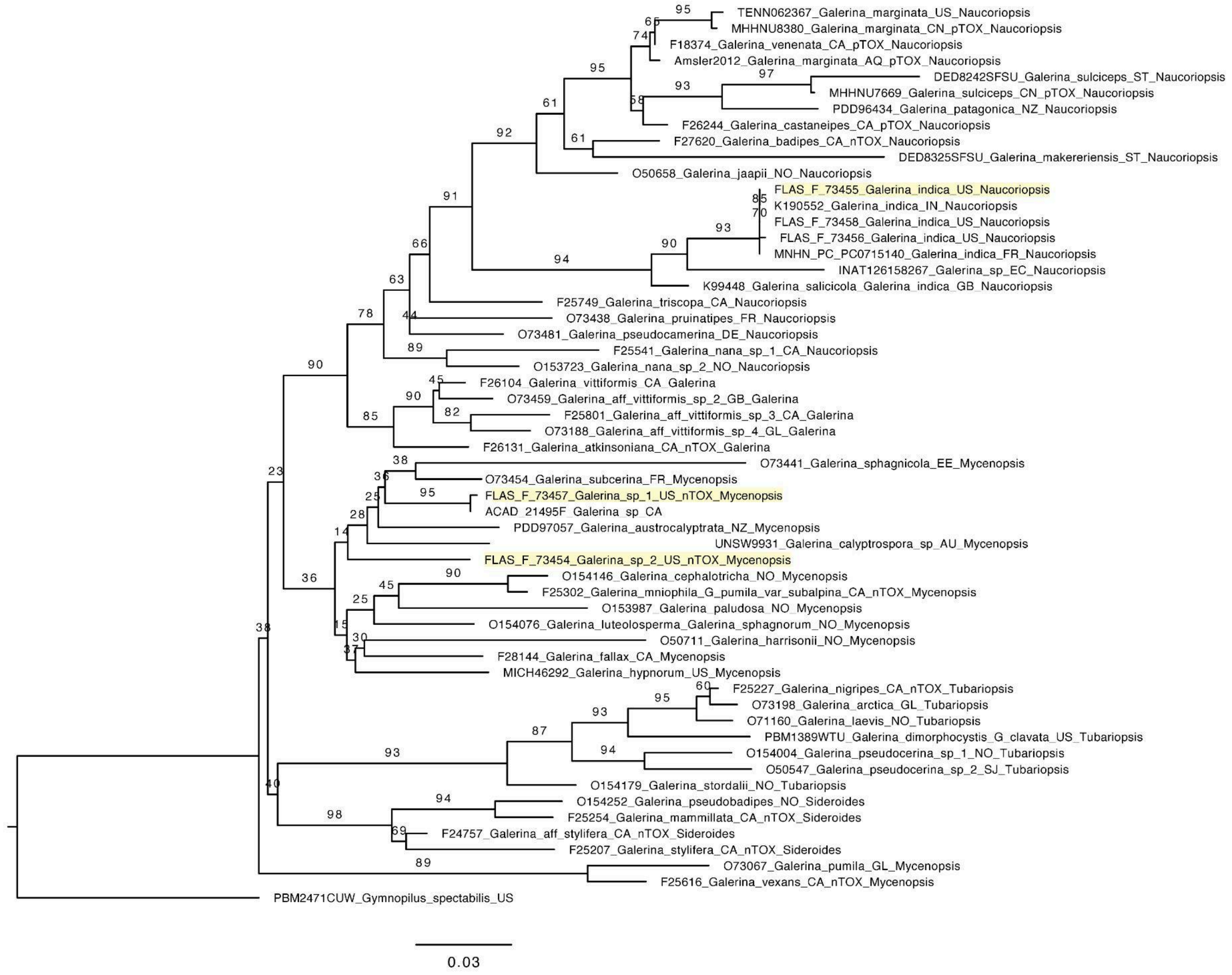


**Figure S1.** Maximum-likelihood phylogeny of *Galerina* inferred from ITS sequences alone (56 taxa, 606 bp; ITS1+ITS2 partition: TVM+F+G4, 5.8S partition: JC) using IQ-TREE 2.4.0. Branch labels indicate ultrafast bootstrap support (1,000 replicates with gene-site resampling). *Gymnopilus spectabilis* is used as an outgroup. FLAS voucher representatives of the three species analyzed as a part of this study are highlighted. Voucher metadata is available in table S2.

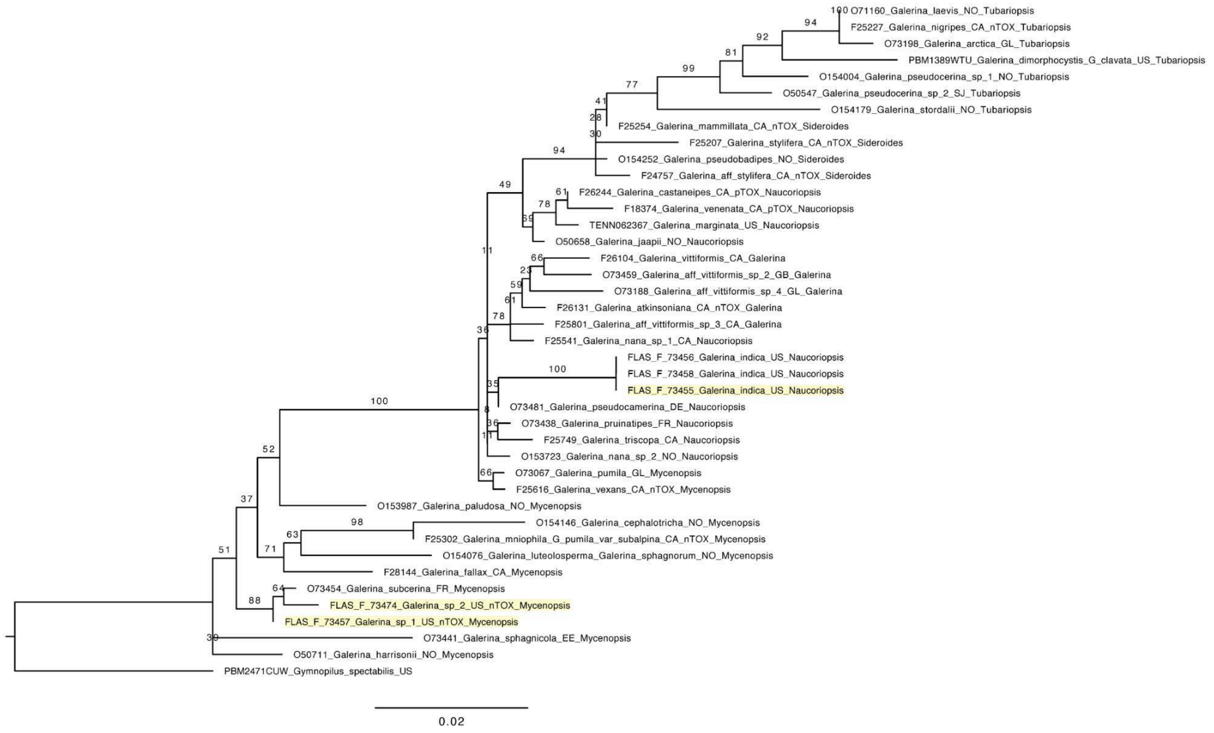


**Figure S2.** Maximum-likelihood phylogeny of *Galerina* inferred from 28S sequences alone (41 taxa, 697 bp; TPM3+R3) using IQ-TREE 2.4.0. Branch labels indicate ultrafast bootstrap support (1,000 replicates). *Gymnopilus subspectabilis* is used as an outgroup. FLAS voucher representatives of the three species analyzed as a part of this study are highlighted. Voucher metadata is available in table S2.

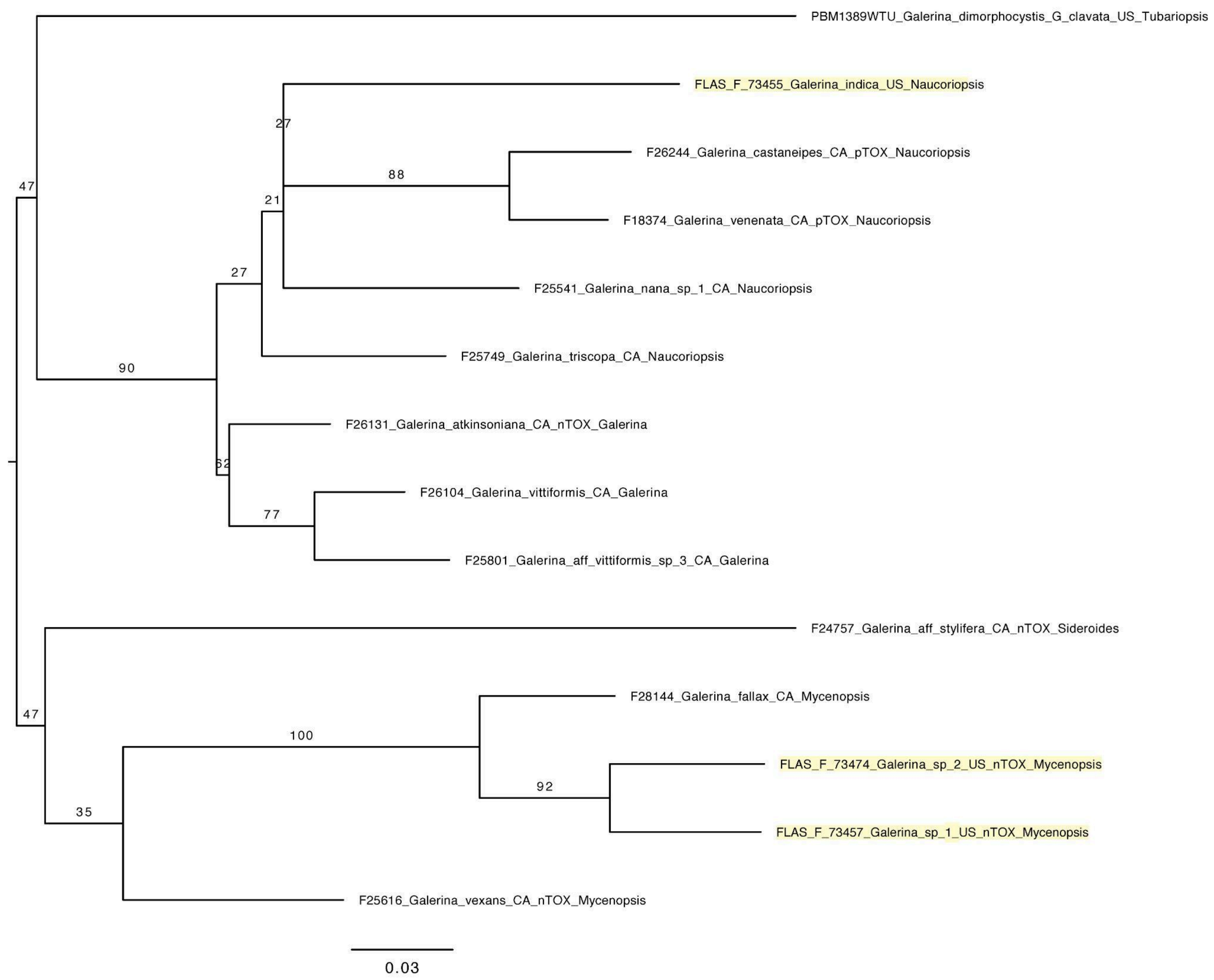


**Figure S3.** Maximum-likelihood phylogeny of *Galerina* inferred from RPB2 sequences alone (14 taxa, 678 bp; codon position 1: TN+F+G4, position 2: JC+I, position 3: HKY+F+G4) using IQ-TREE 2.4.0. Branch labels indicate ultrafast bootstrap support (1,000 replicates with gene-site resampling). The tree is midpoint rooted. FLAS voucher representatives of the three species analyzed as a part of this study are highlighted. Voucher metadata is available in table S2.

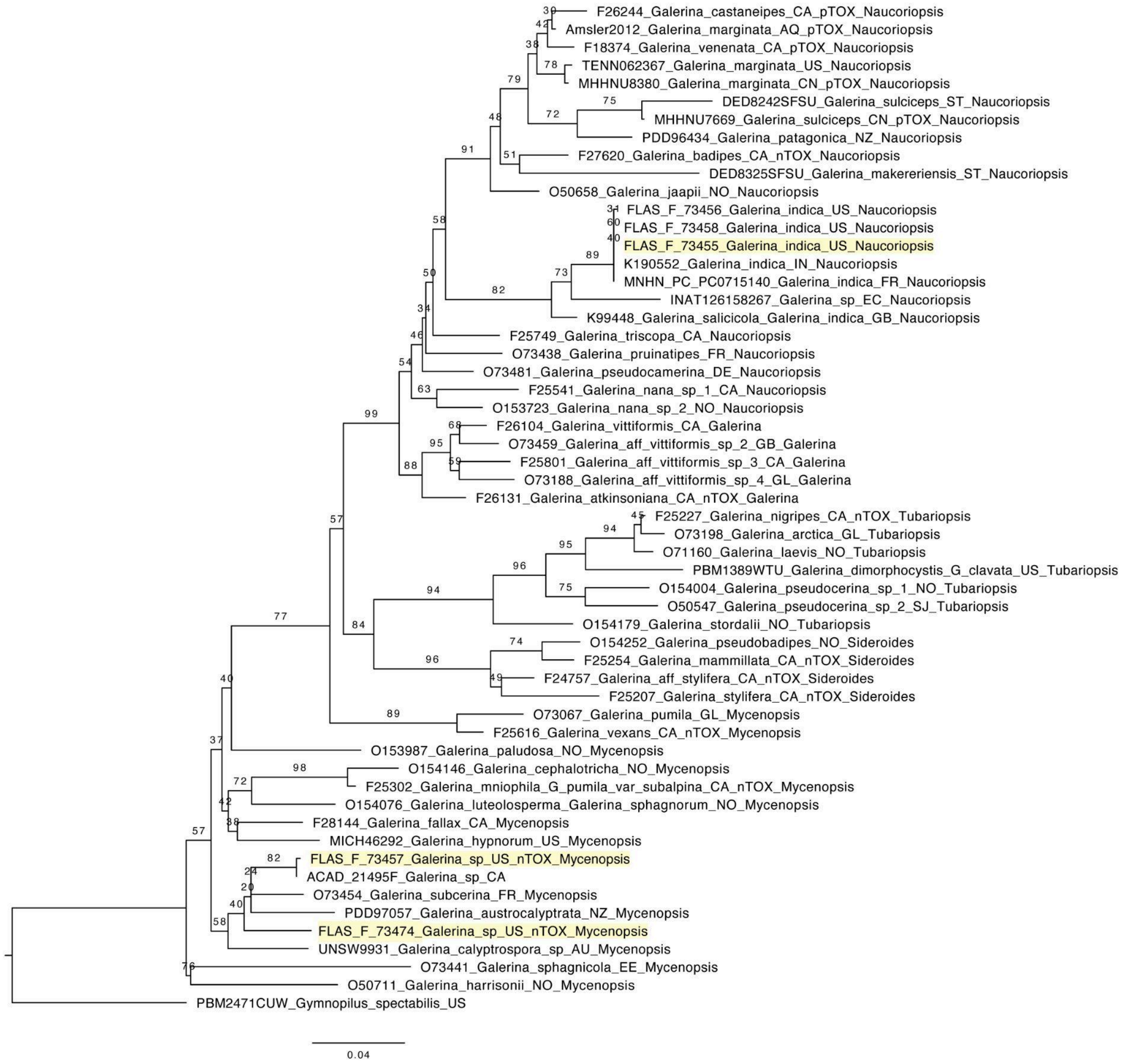


**Figure S4.** Maximum-likelihood phylogeny of *Galerina* inferred from a concatenated alignment of ITS, 28S, and RPB2 (56 taxa, 1,981 bp) using IQ-TREE 2.4.0. This is the same tree from Figure 1, but is shown again here to show all bootstrap values. The matrix was partitioned into seven candidate blocks (ITS1, 5.8S, ITS2, 28S, and RPB2 codon positions 1, 2, and 3) and consolidated by ModelFinder Plus into three final partitions: ITS1+ITS2 (TVM+F+G4), 5.8S+28S+RPB2 codon positions 1+2 (TN+F+I+G4), and RPB2 codon position 3 (HKY+F+G4). Branch labels indicate ultrafast bootstrap support (1,000 replicates with NNI optimization and gene-site resampling). *Gymnopilus subspectabilis* is used as an outgroup. Voucher metadata is available in table S2.

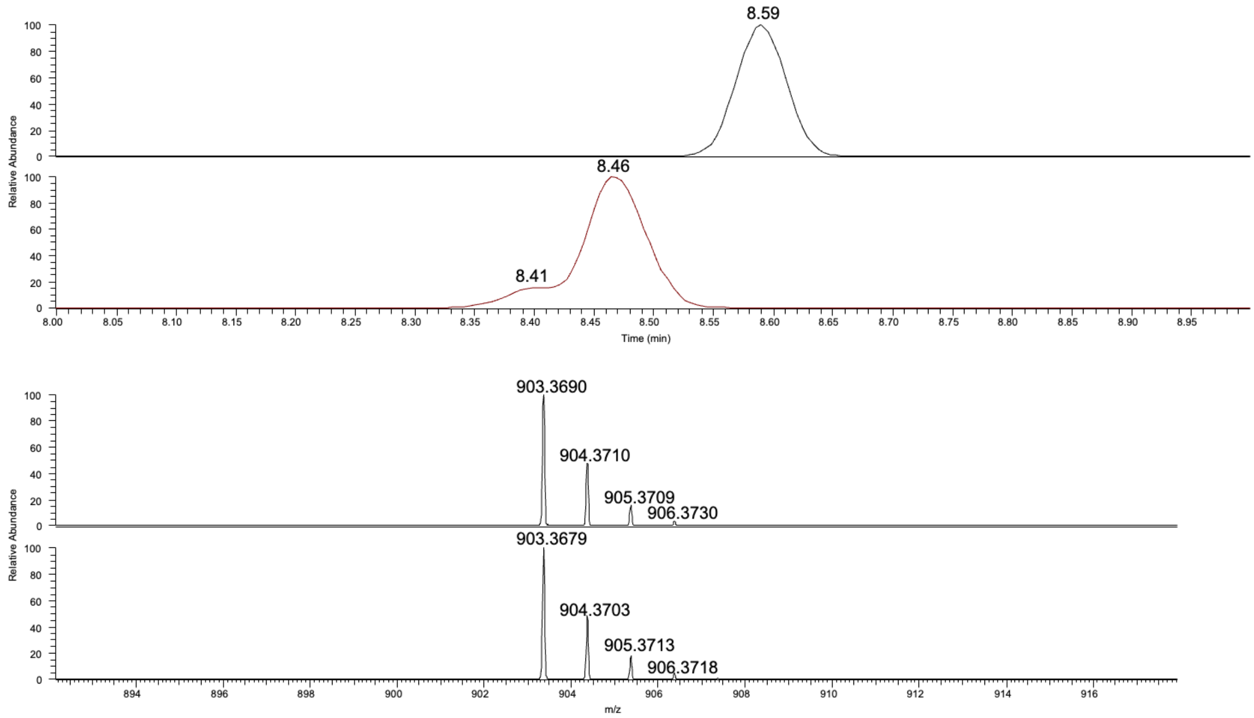


**Figure S5.** EIC of α-proamanitin (*m/z* 903.3525-903.3716) in *Galerina indica* FLAS-F-73455 and for γ-amanitin (red) with their respective extracted mass spectra below.

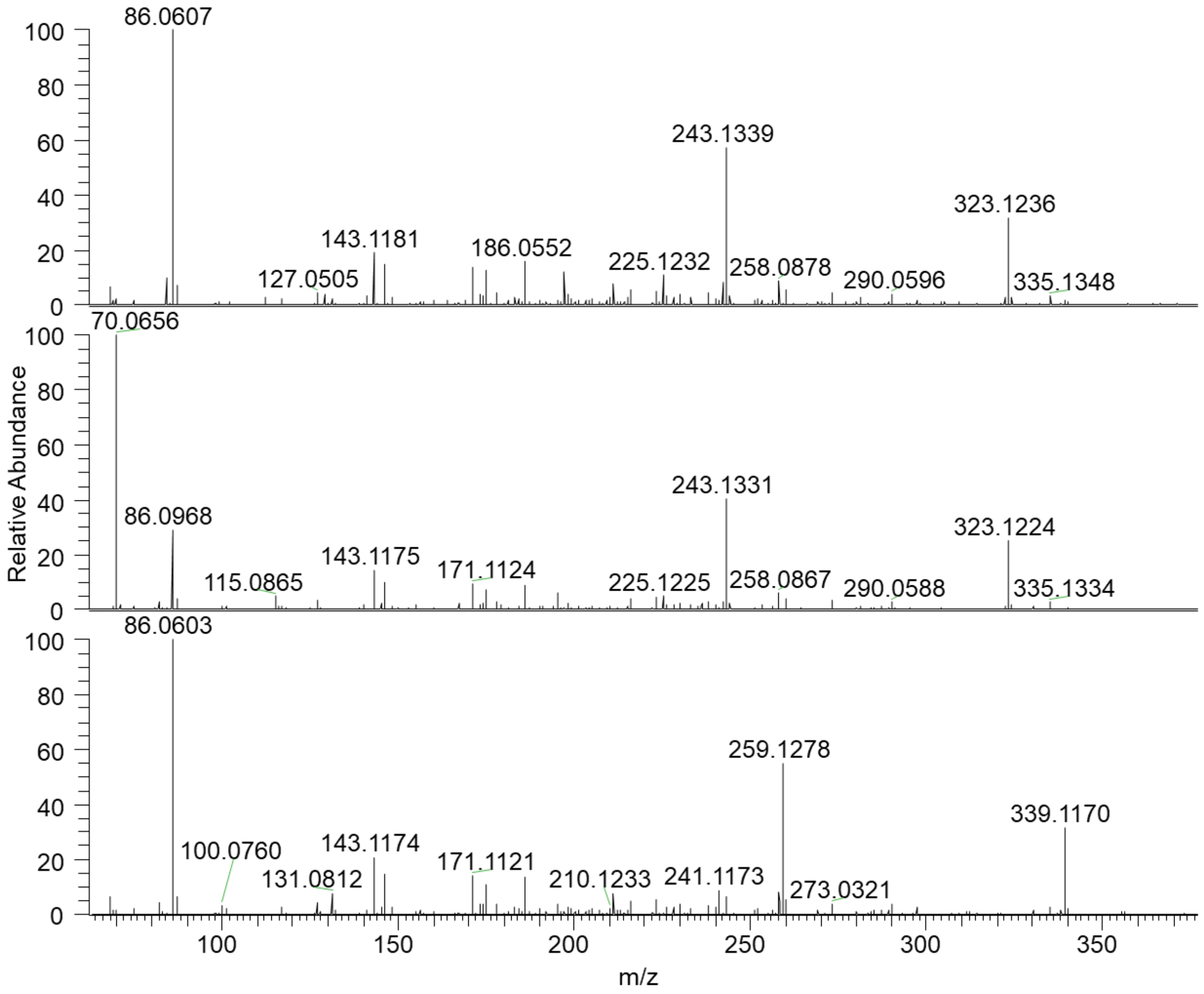


**Figure S6**. Averaged HCD MS/MS spectra of γ-amanitin (top), α-proamanitin (middle), and α-amanitin (bottom) at 35 NCE.

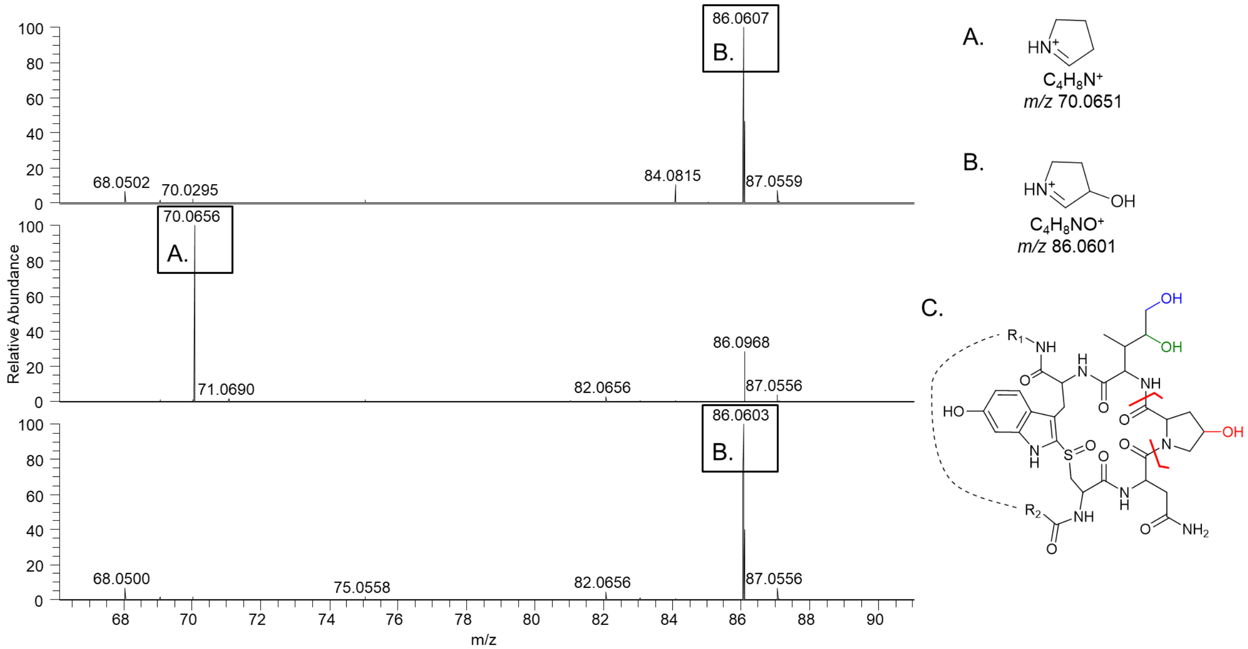


**Figure S7.** Averaged HCD MS/MS spectra of γ-amanitin (top), α-proamanitin (middle), and α-amanitin (bottom) at 35 NCE. The x-axis is zoomed in on the *m/z* range of interest for the proposed proline and hydroxyproline product ion. Corresponding proposed product ion structures are depicted by A, and B. *m/z* 86.0969 is an isobaric interference with ion B. Ion A and B are assumed to form after a secondary loss of CO from the ion that would form from bond cleavages shown in C, for each respective precursor. An example to depict where this residue resides in the peptide is shown on α-amanitin in C. A dotted line connecting $R_1$ and $R_2$ on α-amanitin is used to allow for a more relaxed geometry of the areas of interest.

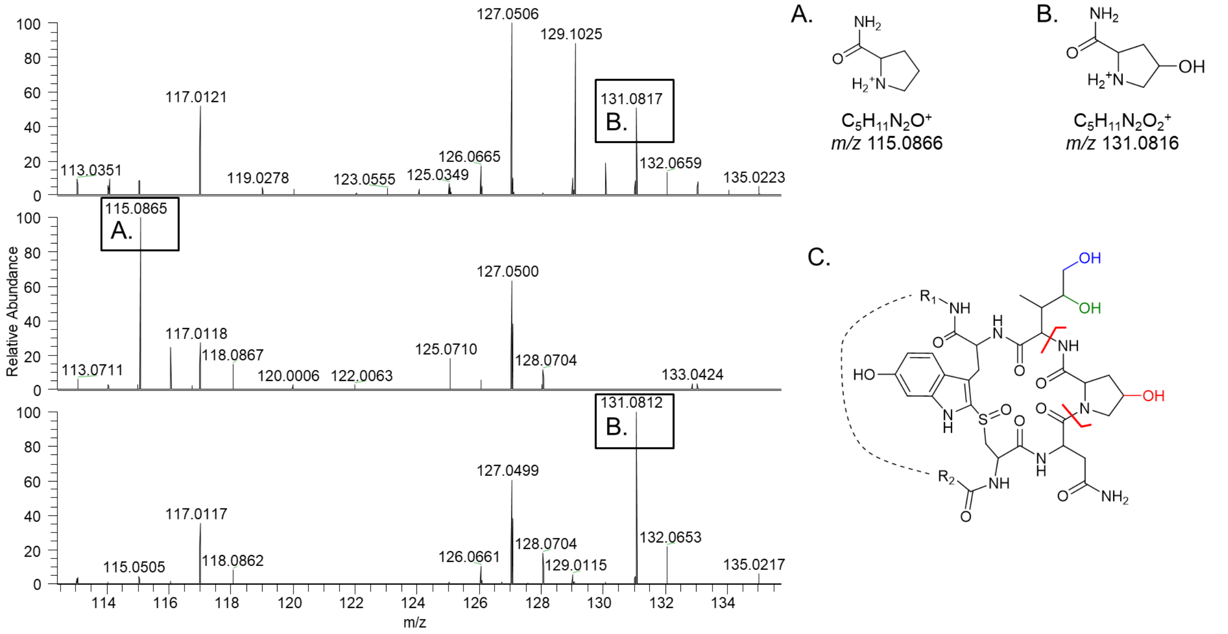


**Figure S8.** Averaged HCD MS/MS spectra of γ-amanitin (top), α-proamanitin (middle), and α-amanitin (bottom) at 35 NCE. The x-axis is zoomed in on the *m/z* range of interest for the proposed proline and hydroxyproline product ion. Corresponding proposed product ion structures are depicted by A, and B. An example of the bond cleavages proposed to form ions A and B from their respective precursor are depicted on α-amanitin shown in C. A dotted line connecting $R_1$ and $R_2$ on α-amanitin is used to allow for a more relaxed geometry of the areas of interest.

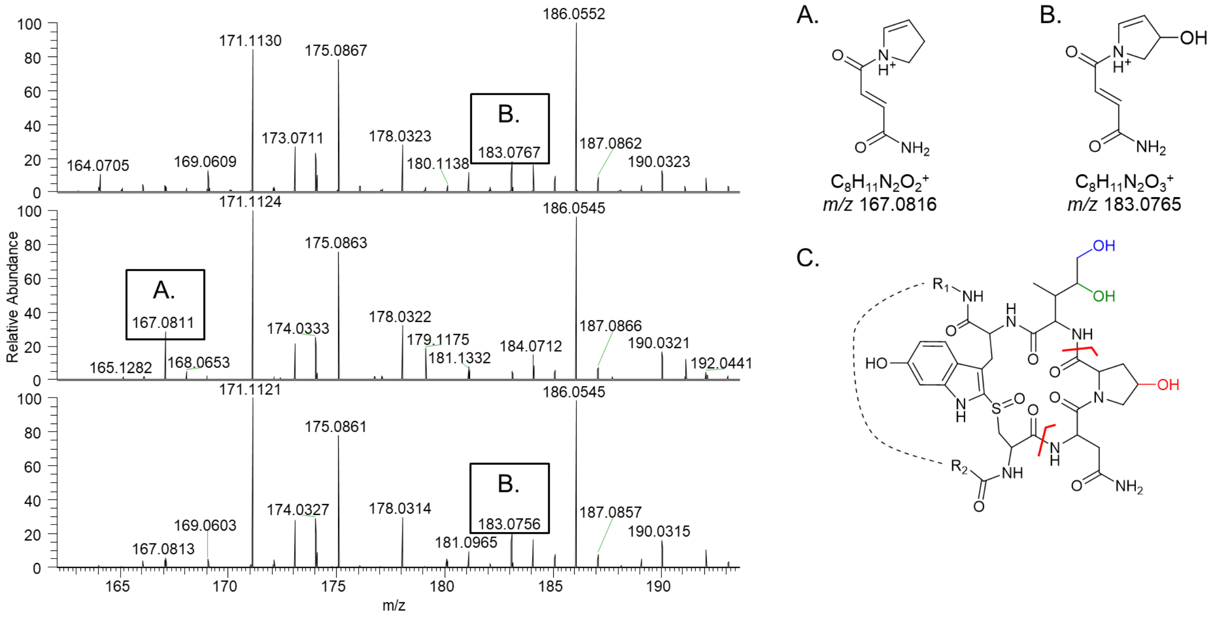


**Figure S9.** Averaged HCD MS/MS spectra of γ-amanitin (top), α-proamanitin (middle), and α-amanitin (bottom) at 35 NCE. The x-axis is zoomed in on the *m/z* range of interest for the proposed proline and hydroxyproline asparagine product ion. Corresponding proposed product ion structures are assumed to form after secondary loss of CO and $NH_3$, and are depicted by A, and B. An example of the bond cleavages proposed to form ions A and B from their respective precursor are depicted on α-amanitin shown in C. A dotted line connecting $R_1$ and $R_2$ on α-amanitin is used to allow for a more relaxed geometry of the areas of interest.

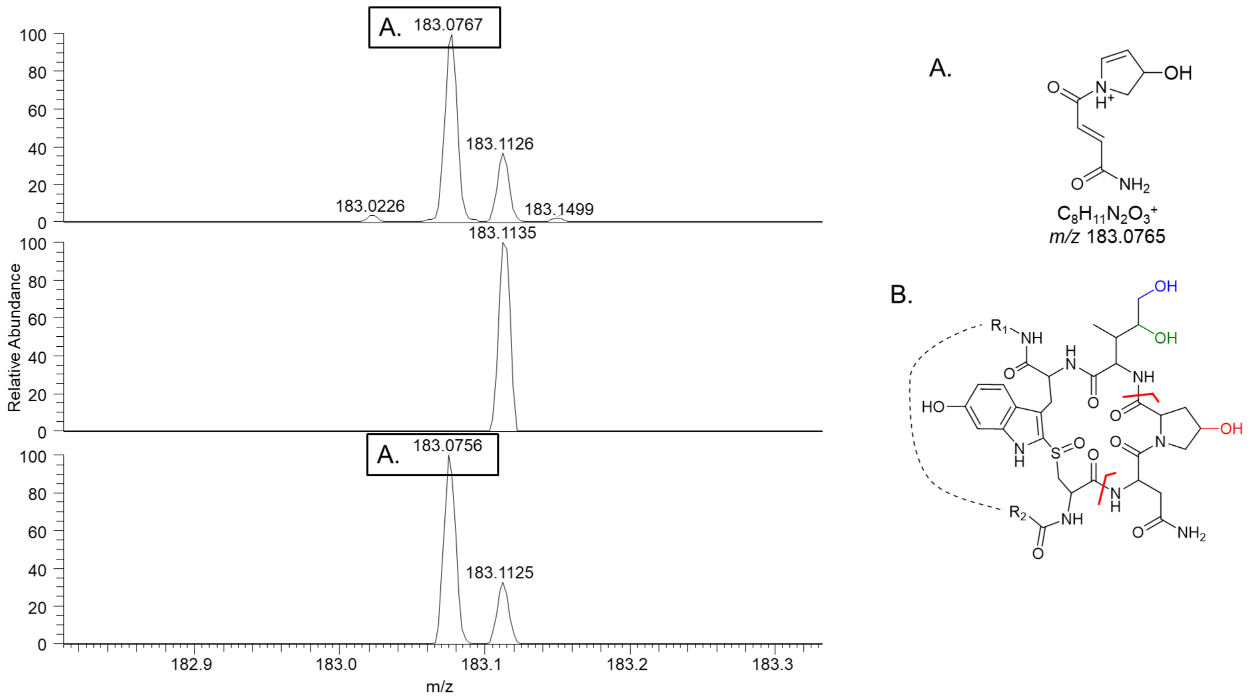


**Figure S10.** Averaged HCD MS/MS spectra of γ-amanitin (top), α-proamanitin (middle), and α-amanitin (bottom) at 35 NCE. The x-axis is zoomed in on the *m/z* range of interest for the proposed hydroxyproline asparagine product ion and the isobaric interference. The corresponding proposed product ion structure is assumed to form after secondary loss of CO and $NH_3$, and is depicted by A. An example of the bond cleavages proposed to form the ion from the respective precursor is depicted on α-amanitin shown in B. A dotted line connecting $R_1$ and $R_2$ on α-amanitin is used to allow for a more relaxed geometry of the areas of interest.

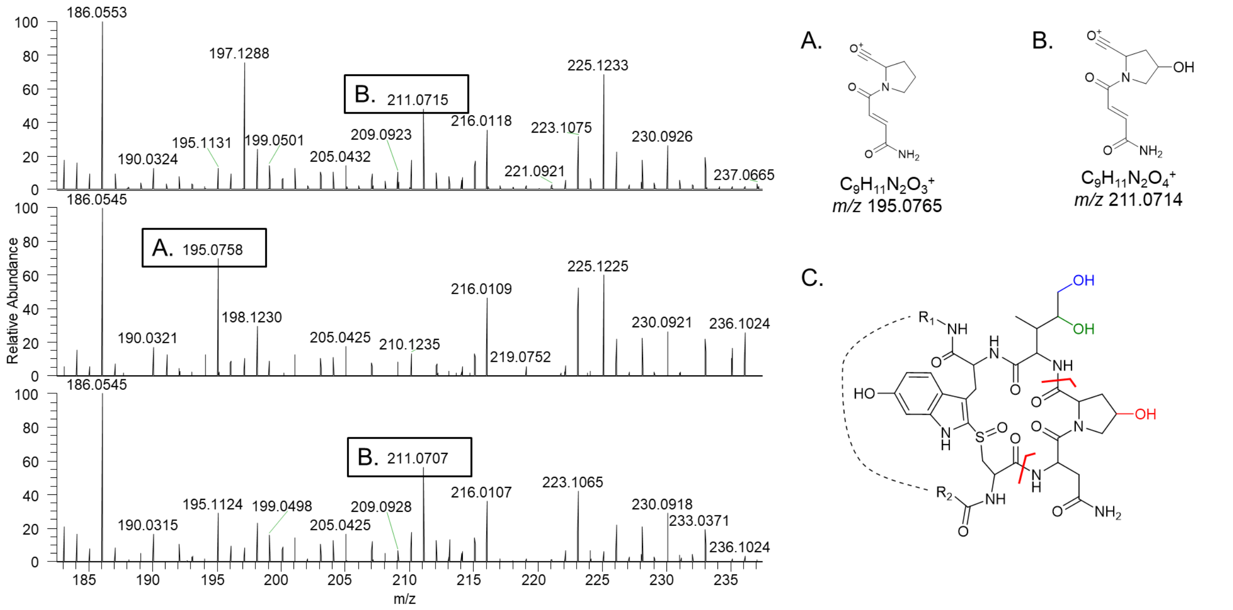


**Figure S11.** Averaged HCD MS/MS spectra of γ-amanitin (top), α-proamanitin (middle), and α-amanitin (bottom) at 35 NCE. The x-axis is zoomed in on the *m/z* range of interest for the proposed proline and hydroxyproline asparagine product ion. Corresponding proposed product ion structures are assumed to form after secondary loss of NH3, and are depicted by A, and B. An example of the bond cleavages proposed to form ions A and B from their respective precursor are depicted on α-amanitin shown in C. A dotted line connecting $R_1$ and $R_2$ on α-amanitin is used to allow for a more relaxed geometry of the areas of interest.

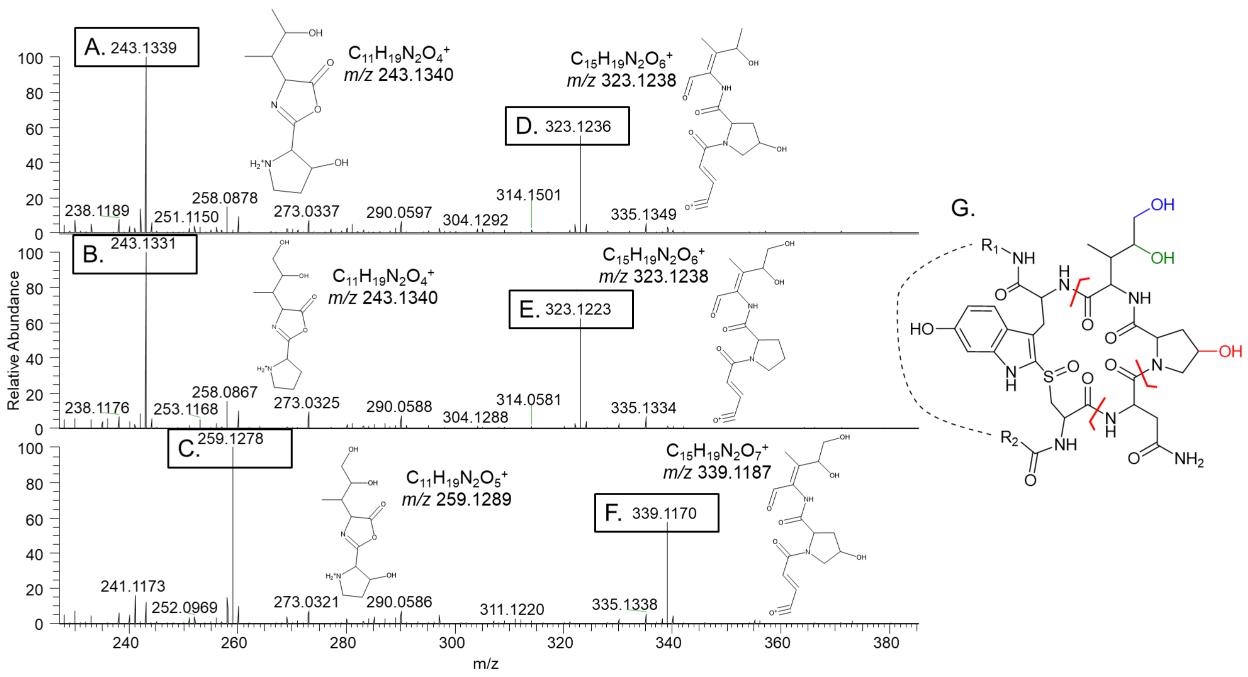


**Figure S12.** Averaged HCD MS/MS spectra of γ-amanitin (top), α-proamanitin (middle), and α-amanitin (bottom) at 35 NCE. The x-axis is zoomed in on the *m/z* range of interest for the two largest product ions which are isomeric and of significant intensity. Corresponding proposed product ion structures are assumed to form after secondary losses and or rearrangements, and are depicted by A-F. An example of the bond cleavages proposed to form ions A-F from their respective precursor are depicted on α-amanitin shown in G. A dotted line connecting $R_1$ and $R_2$ on α-amanitin is used to allow for a more relaxed geometry of the areas of interest.

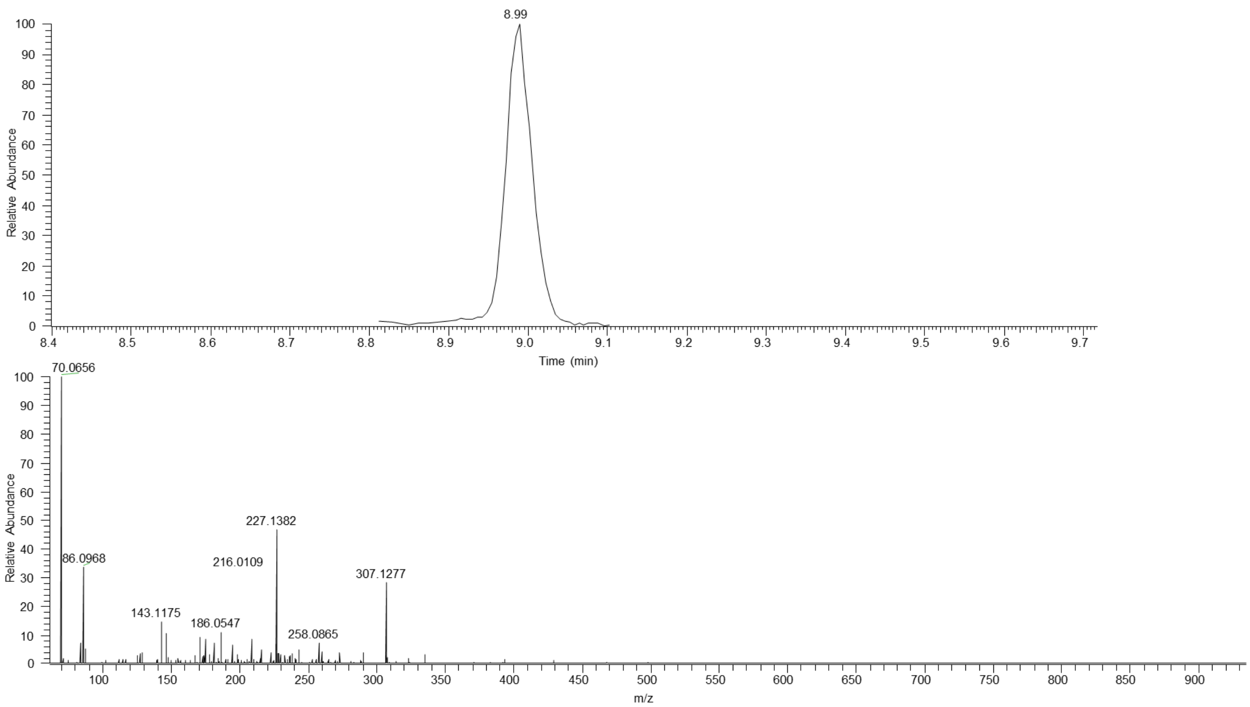


**Figure S13.** EIC of MS/MS scans corresponding to *m/z* 887.370 in *Galerina indica* (FLAS-F-73455). An averaged MS/MS spectrum under HCD NCE 35 is shown.

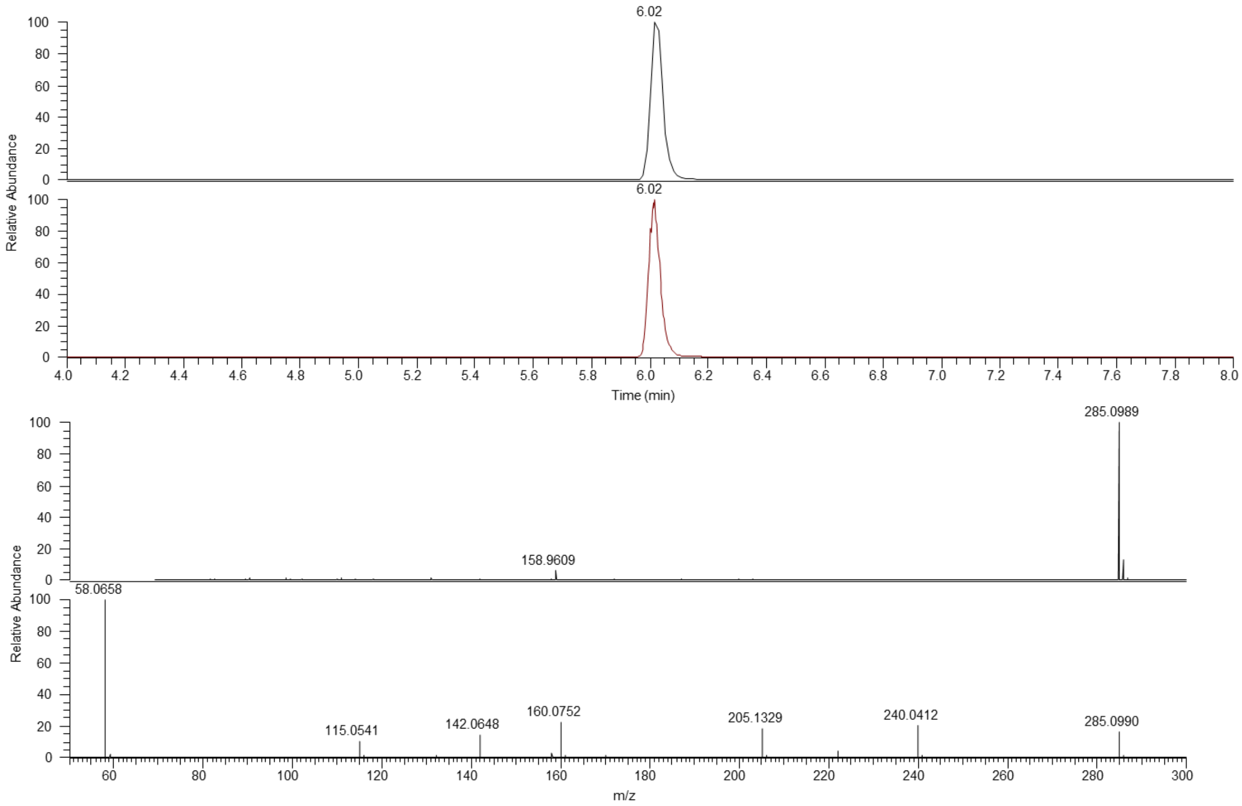


**Figure S14.** EIC of *m/z* 285.0970-285.1028, corresponding to psilocybin $[M+H]^+$, in *Galerina indica* (FLAS-F-73455) (black) and *Galerina* sp. 1 (FLAS-F-73457) (red). A representative extracted mass spectra and MS/MS spectrum under HCD NCE 35 is shown for *Galerina indica* (FLAS-F-73455).

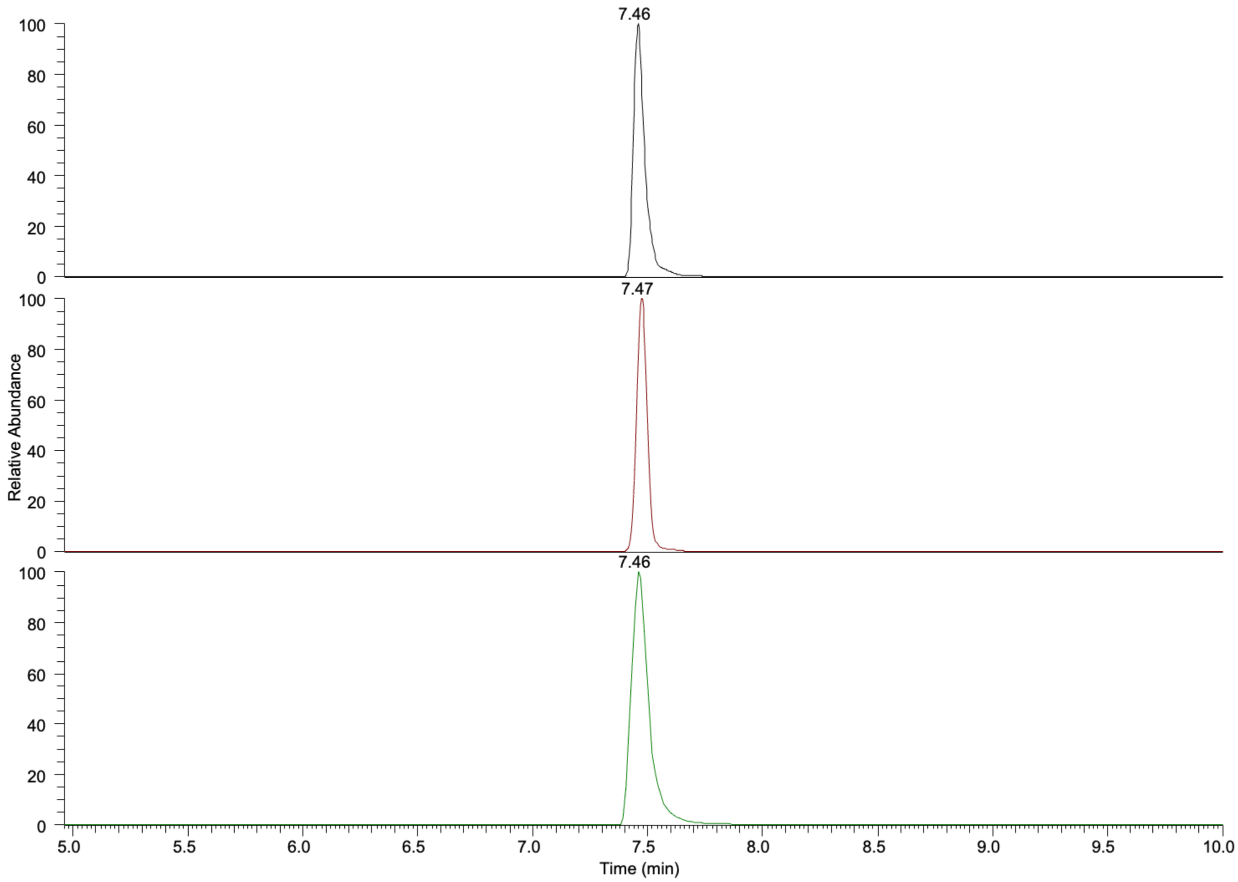


**Figure S15.** EIC of *m/z* 205.1314-205.1356, corresponding to psilocin $[M+H]^+$, from a certified reference material at 1 μM (middle), *Galerina indica* (FLAS-F-73455) (bottom) and *Galerina* sp. 1 (FLAS-F-73457) (bottom).

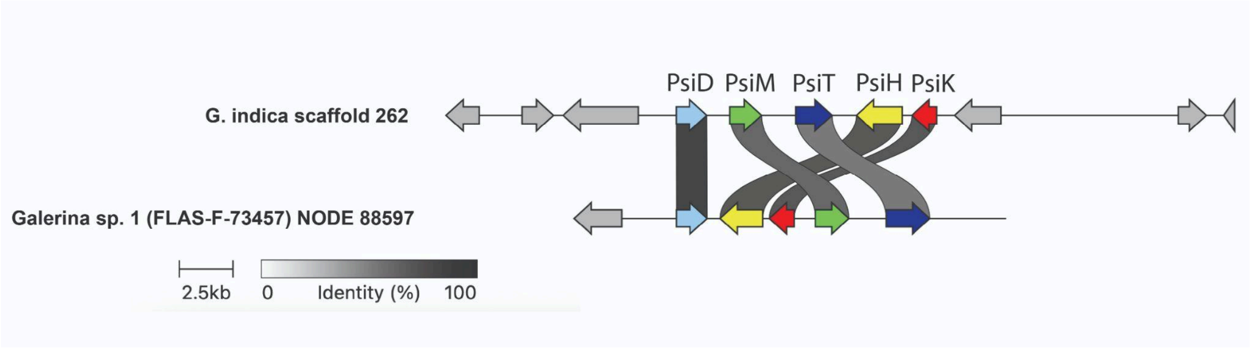


**Figure S16.** Synteny comparison of *Galerina indica* (FLAS-F-73455) and *Galerina* sp. 1 (FLAS-F-73457) *Psi* gene loci. Genes are colored by identity. The *G. indica* locus is from BioSample SAMN60278622 and *Galerina* sp. 1 (FLAS-F-73457) is from biosample SAMN60278690**.**

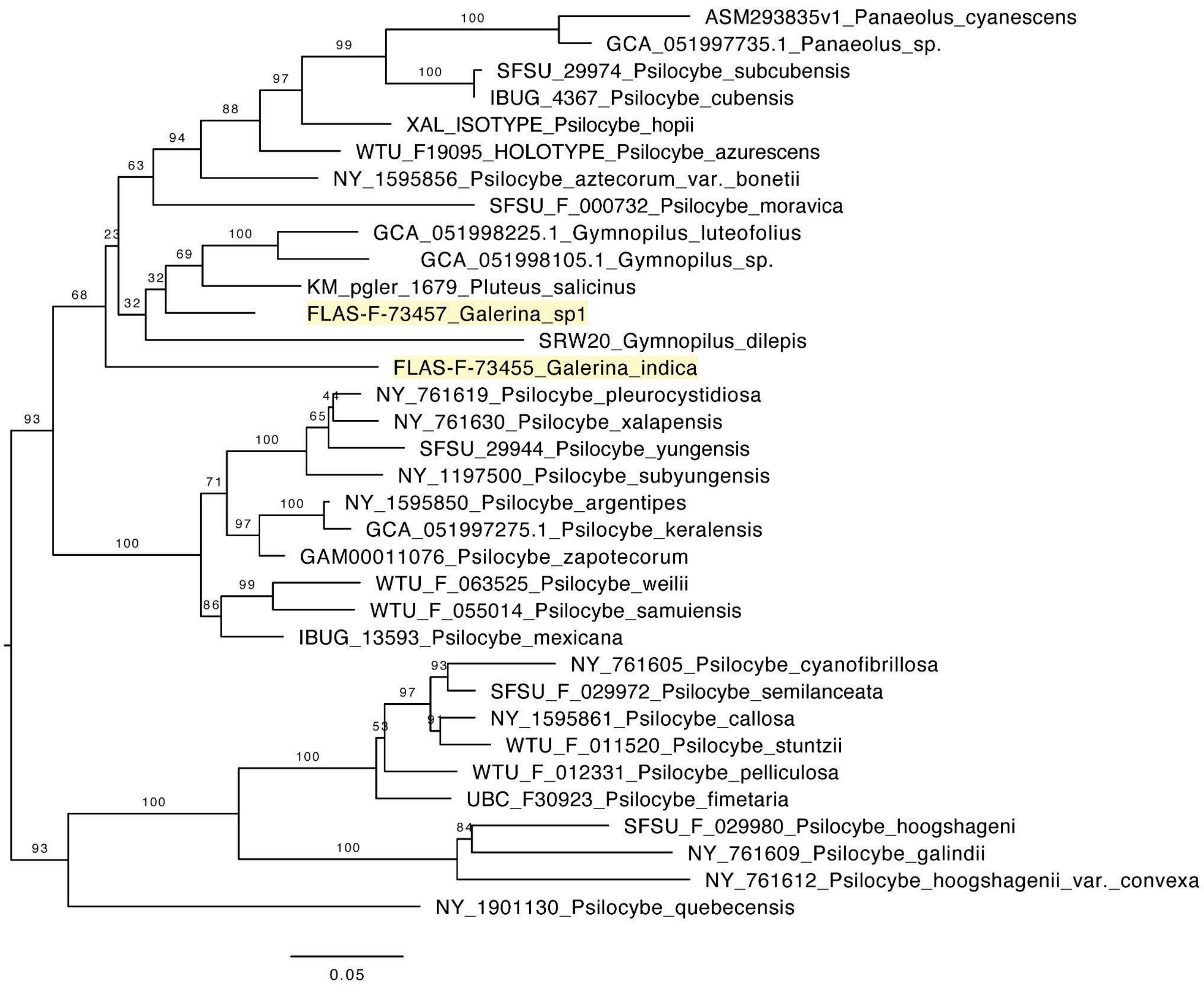


**Figure S17.** Maximum likelihood phylogeny of *PsiH* (34 taxa, 472 amino acid positions) inferred in RAxML v8.2.10 under the JTTDCMUT+Γ substitution model. Node values indicate bootstrap support from 500 rapid bootstrap replicates. The tree is midpoint-rooted. Voucher metadata are available in table S4.

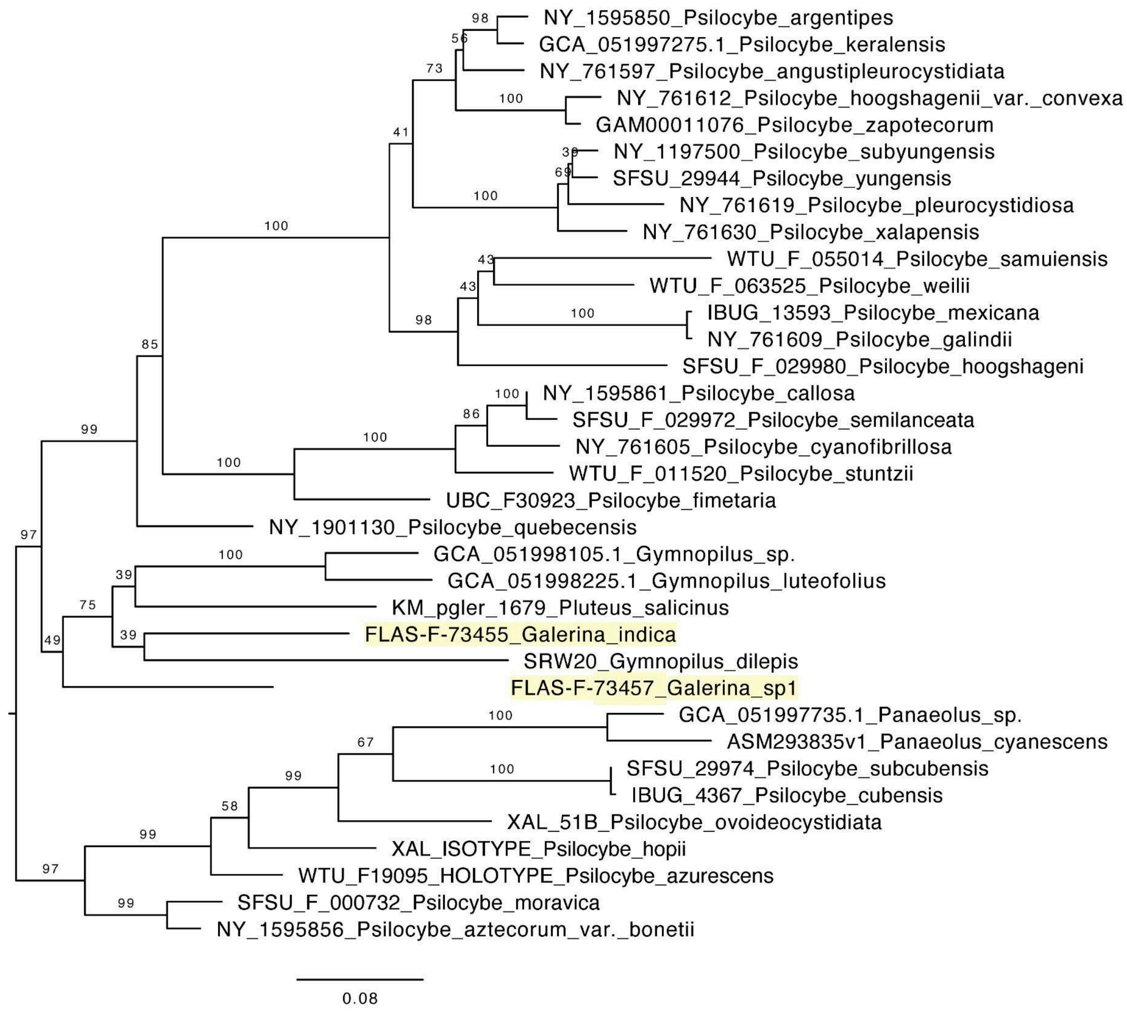


**Figure S18.** Maximum likelihood phylogeny of *PsiK* (35 taxa, 351 amino acid positions) inferred in RAxML v8.2.10 under the JTT+Γ substitution model. Node values indicate bootstrap support from 500 rapid bootstrap replicates.The tree is midpoint-rooted. Voucher metadata are available in table S4.

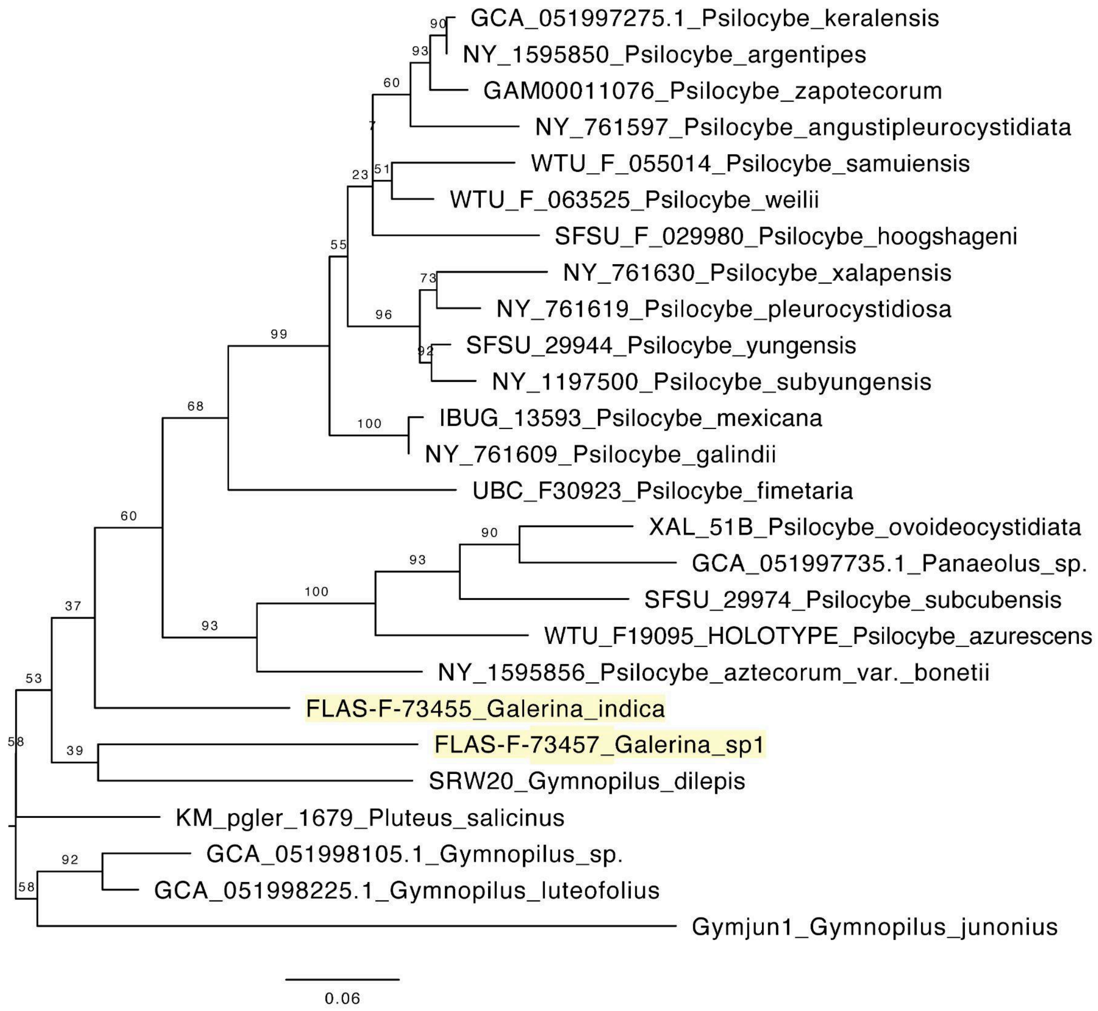


**Figure S19.** Maximum likelihood phylogeny of *PsiM* (26 taxa, 294 amino acid positions) inferred in RAxML v8.2.10 under the CPREV+Γ substitution model. Node values indicate bootstrap support from 500 rapid bootstrap replicates. The tree is midpoint-rooted. Voucher metadata is available in table S4.

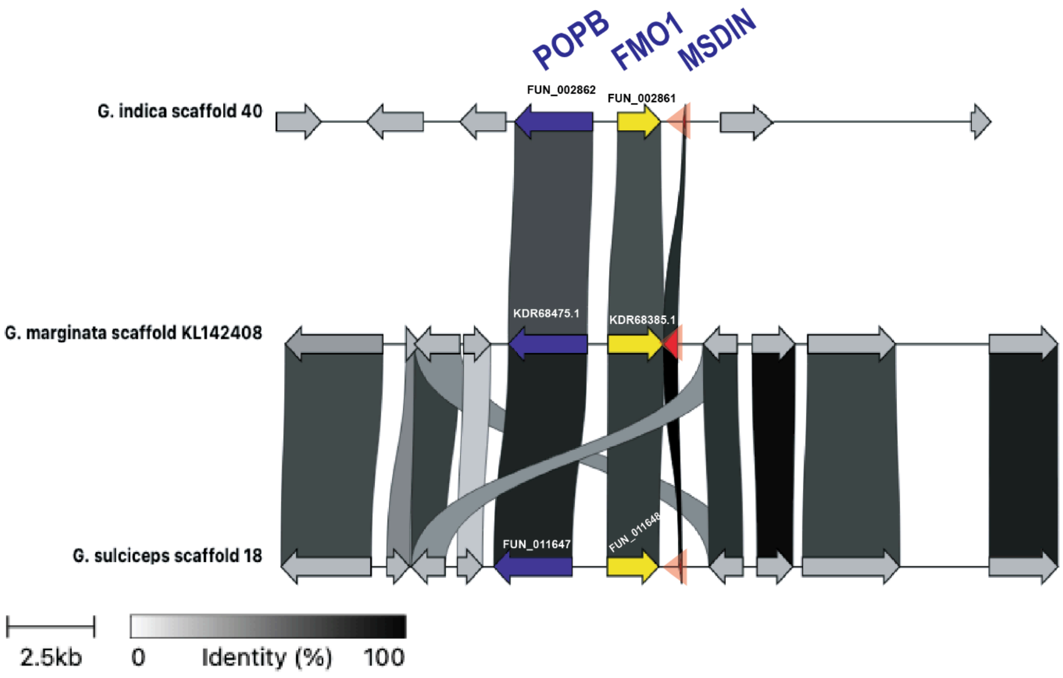


**Figure S20.** Amatoxin biosynthesis gene cluster Synteny comparison. Taxa represented include *Galerina indica* (FLAS-F-73455 scaffold 40), *G. marginata* (Galma1 CBS339.88 scaffold KL142408), *G. sulciceps* (ZP082096 scaffold 18 ). Genes not known to be implicated in amatoxin biosynthesis are shown in grey. The MSDIN genes are highlighted with a translucent red arrow for visibility. The *G. indica* MSDIN copy shown here is the inferred functional copy shown in Fig 2. The *G. marginata* MSDIN copy shown here on scaffold KL142408 is orthologous to the Gm1b(AMA-1) copy shown in Fig 2. The G. *sulciceps* MSDIN copy shown here is orthologous to the Gs1b(AMA-1) functional copy shown in Fig 2.

### Tables

**Table S1.** (Separate file) Voucher metadata for *Galerina* taxa and others analyzed as a part of this study.

**Table S2.** (Separate file) Voucher metadata for ITS, 28S, and *RPB2* sequences and information on toxicity for vouchers analyzed in *Galerina* organismal phylogeny (Figure 1).

**Table S3.** Genome assembly metrics, including busco scores and assembly size, of the *Galerina* specimens sequenced as a part of this study.

| **Voucher:** | ***Galerina* sp. 1 (FLAS-F-73457)** | ***Galerina indica* (FLAS-F-73455)** |
|---|---|---|
| **ASSEMBLY STATISTICS** | | |
| **Technology (see methods)** | Illumina only | Hybrid (NANOPORE + illumina polish) |
| **Assembly Length (Mb)** | 63.2 | 61.65 |
| **Number of Contigs** | 38,899 | 1,496 |
| **N50 (bp)** | 2,415 | 90,626 |
| **Max Contig (bp)** | 1,212,504 | 630,902 |
| **GC Content (%)** | 47.89% | 47.24 |
| **BUSCO (agaricales_odb10, n=3870)** | | |
| **Complete (C%)** | 68.1% | 98.2 |
| **Single-copy (S%)** | 67.2% | 93.9 |
| **Duplicated (D%)** | 0.9% | 4.3 |
| **Fragmented (F%)** | 11.1% | 0.5 |
| **Missing (M%)** | 20.8% | 1.3 |
| **Tool Version** | BUSCO 5.3.0 | BUSCO 5.8.3 |
| **Gene Predictor** | metaeuk | miniprot 0.14-r265 |
| **COMPLEASM (agaricales_odb12, n=3372)** | | |
| **Single-copy (S%)** | 65.36% | 92.08 |
| **Duplicated (D%)** | 0.36% | 3.59 |
| **Fragmented (F%)** | 19.87% | 0.89 |
| **Incomplete (I%)** | 0.21% | 0 |
| **Missing (M%)** | 14.21% | 3.44 |
| **Tool Version** | Compleasm 0.2.7 | Compleasm 0.2.7 |

**Table S4.** Voucher metadata for various *Psi*-containing taxa used in the Psi protein phylogeny shown in Figure 3.

| Species | Voucher/Strain/ Accession | Genes available | Reference |
|---|---|---|---|
| *Psilocybe angustipleurocystidiata* | NY761597 | PsiD/PsiK/PsiM | Bradshaw et al. (2024) |
| *Psilocybe argentipes* | NY1595850 | PsiD/PsiH/PsiK/PsiM | Bradshaw et al. (2024) |
| *Psilocybe aztecorum* var. *bonetii* | NY1595856 | PsiD/PsiH/PsiK/PsiM | Bradshaw et al. (2024) |
| *Psilocybe azurescens* | WTUF19095 | PsiD/PsiH/PsiK/PsiM | Bradshaw et al. (2024) |
| *Psilocybe callosa* | NY1595861 | PsiD/PsiH/PsiK | Bradshaw et al. (2024) |
| *Psilocybe cubensis* | IBUG4367 | PsiD/PsiH/PsiK | Bradshaw et al. (2024) |
| *Psilocybe cyanofibrillosa* | NY761605 | PsiD/PsiH/PsiK | Bradshaw et al. (2024) |
| *Psilocybe fimetaria* | UBC F30923 | PsiD/PsiH/PsiK/PsiM | Bradshaw et al. (2024) |
| *Psilocybe galindii* | NY761609 | PsiD/PsiH/PsiK/PsiM | Bradshaw et al. (2024) |
| *Psilocybe hoogshageni* | SFSUF029980 | PsiD/PsiH/PsiK/PsiM | Bradshaw et al. (2024) |
| *Psilocybe hoogshagenii* var. *convexa* | NY761612 | PsiD/PsiH/PsiK | Bradshaw et al. (2024) |
| *Psilocybe hopii* | XAL ISOTYPE | PsiD/PsiH/PsiK | Bradshaw et al. (2024) |
| *Psilocybe mexicana* | IBUG13593 | PsiD/PsiH/PsiK/PsiM | Bradshaw et al. (2024) |
| *Psilocybe moravica* | SFSU F000732 | PsiD/PsiH/PsiK | Bradshaw et al. (2024) |
| *Psilocybe ovoideocystidiata* | XAL51B | PsiD/PsiK/PsiM | Bradshaw et al. (2024) |
| *Psilocybe pelliculosa* | WTU F012331 | PsiD/PsiH | Bradshaw et al. (2024) |
| *Psilocybe pleurocystidiosa* | NY761619 | PsiD/PsiH/PsiK/PsiM | Bradshaw et al. (2024) |
| *Psilocybe quebecensis* | NY1901130 | PsiD/PsiH/PsiK | Bradshaw et al. (2024) |
| *Psilocybe samuiensis* | WTU F055014 | PsiD/PsiH/PsiK/PsiM | Bradshaw et al. (2024) |
| *Psilocybe semilanceata* | SFSU F029972 | PsiD/PsiH/PsiK | Bradshaw et al. (2024) |
| *Psilocybe stuntzii* | WTU F011520 | PsiD/PsiH/PsiK | Bradshaw et al. (2024) |
| *Psilocybe subcubensis* | SFSU29974 | PsiD/PsiH/PsiK/PsiM | Bradshaw et al. (2024) |
| *Psilocybe subyungensis* | NY1197500 | PsiD/PsiH/PsiK/PsiM | Bradshaw et al. (2024) |
| *Psilocybe weilii* | WTU F063525 | PsiD/PsiH/PsiK/PsiM | Bradshaw et al. (2024) |
| *Psilocybe xalapensis* | NY761630 | PsiD/PsiH/PsiK/PsiM | Bradshaw et al. (2024) |
| *Psilocybe yungensis* | SFSU F29944 | PsiD/PsiH/PsiK/PsiM | Bradshaw et al. (2024) |
| *Psilocybe zapotecorum* | GAM00011076 | PsiD/PsiH/PsiK/PsiM | Bradshaw et al. (2024) |
| *Panaeolus cyanescens* | ASM293835v1 | PsiD/PsiH/PsiK | Bradshaw et al. (2024) |
| *Pholiotina smithii* | 2634DeC | PsiD | Bradshaw et al. (2024) |
| *Pluteus salicinus* | KM pgler1679 | PsiD/PsiH/PsiK/PsiM | Bradshaw et al. (2024) |
| *Pluteus albostipitatus* | KM54312 | PsiD | Bradshaw et al. (2024) |
| *Gymnopilus chrysopellus* | Gymch1 PR-1187 | PsiD | Grigoriev et al. (2014) |
| *Gymnopilus dilepis* | SRW20 | PsiD/PsiH/PsiK/PsiM | Reynolds et al. (2018) |
| *Gymnopilus junonius* | Gymjun1 | PsiD, PsM | Ruiz-Dueñas et al. (2021) |
| *Galerina indica* | FLAS-F-73455 | PsiD/PsiH/PsiK/PsiM | This publication |

| | | | |
|---|---|---|---|
| *Galerina sp. 1* | FLAS-F-73457 | PsiD/PsiH/PsiK/PsiM | This publication |
| *Psilocybe keralensis* | GCA_05199727 5.1 | PsiD/PsiH/PsiK/PsiM | Liu et al. (2025) |
| *Panaeolus* sp. | GCA_05199773 5.1 | PsiD/PsiH/PsiK/PsiM | Liu et al. (2025) |
| *Gymnopilus* sp. | GCA_05199810 5.1 | PsiD/PsiH/PsiK/PsiM | Liu et al. (2025) |
| *Gymnopilus luteofolius* | GCA_05199822 5.1 | PsiD/PsiH/PsiK/PsiM | Liu et al. (2025) |

**Table S5.** Voucher metadata for various amatoxin-containing taxa used in the MSDIN protein alignment shown in Figure 2. The putatively nonfunctional copies are marked with a red asterisk as in Figure 2.

| Species | MSDIN Gene identifier | Voucher/strain | NCBI/GenBank | Reference |
|---|---|---|---|---|
| *Galerina indica* | putative functional copy | FLAS-F-73455 | SAMN60278622 | This publication |
| *Galerina indica* | putative nonfunctional copy* | FLAS-F-73455 | SAMN60278622 | This publication |
| *Galerina marginata* | Gm1[b] (alpha-AMA1) | MHHNU 8380 | MN272413 | He et al. (2020) |
| *Galerina marginata* | Gm2[b] (alpha-AMA2) | MHHNU 8380 | MN272414 | He et al. (2020) |
| *Galerina suliceps* | Gs1[b] (alpha-AMA1) | MHHNU 7669 | MN272417 | He et al. (2020) |
| *Galerina suliceps* | Gs2[b] nonfunctional* (alpha-AMA2) | MHHNU 7669 | MN272418 | He et al. (2020) |
| *Amanita subpallidorosea* | Asp2[ab] | MHHNU 8617 | MN264224 | He et al. (2020) |
| *Lepiota venata* | Lv1[b] | MHHNU 31031 | MN272421 | He et al. (2020) |

**Table S6.** Voucher metadata for various amatoxin-containing taxa used in the POPB phylogeny shown in Figure 2.

| Species | Voucher/strain | NCBI/GenBank | Reference |
|---|---|---|---|
| *Hypsizygus marmoreus* | 51987-8 | LUEZ02000233.1 | Choi et al. unpublished |
| *Termitomyces* sp. | J132 | GCA_001263195.1 | Kreuzenbeck et al. (2022) |
| *Lepiota venenata* | MHHNU31031 | MN272424 | He et al. (2020) |
| *Amanita molliuscula* | MHHNU9142 | MN061272 | He et al. (2020) |
| *Amanita exitialis* | MHHNU30937 | KR996717 | He et al. (2020) |
| *Amanita subpallidorosea* | MHHNU8617 | KU601411 | He et al. (2020) |
| *Amanita rimosa* | MHHNU9050 | MN061275 | He et al. (2020) |
| *Amanita fuliginea* | MHHNU9047 | MN061271 | He et al. (2020) |
| *Amanita pallidorosea* | MHHNU31203 | MN061274 | He et al. (2020) |
| *Galerina indica* | FLAS-F-73455 | SAMN60278622 | He et al. (2020) |
| *Galerina marginata* | MHHNU8380 | MN272416 | This publication |
| *Galerina sulciceps* | MHHNU7669 | MN272420 | This publication |
